# Machine learning approaches to identify sleep genes

**DOI:** 10.1101/2021.04.10.439249

**Authors:** Yin Yeng Lee, Mehari Endale, Gang Wu, Marc D Ruben, Lauren J Francey, Andrew R Morris, Natalie Y Choo, Ron C Anafi, David F Smith, Andrew Chuanyin Liu, John B Hogenesch

**Affiliations:** Divisions of Human Genetics and Immunobiology, Department of Pediatrics, Cincinnati Children’s Hospital Medical Center, Cincinnati, OH, 45229, USA; Department of Pharmacology and Systems Physiology, University of Cincinnati College of Medicine, Cincinnati, OH 45229, USA; Department of Physiology and Functional Genomics, University of Florida College of Medicine, Gainesville, FL 32610, USA; Division of Pediatric Otolaryngology-Head and Neck Surgery, Cincinnati Children’s Hospital Medical Center, Cincinnati, OH 45229, USA; Division of Pulmonary Medicine and the Sleep Center, Cincinnati Children’s Hospital Medical Center, Cincinnati, OH 45229, USA; The Center for Circadian Medicine, Cincinnati Children’s Hospital Medical Center, Cincinnati, OH 45229, USA; Department of Medicine, Chronobiology and Sleep Institute, Perelman School of Medicine, University of Pennsylvania, Philadelphia, PA 19104, USA

## Abstract

Genetics impacts sleep, yet, the molecular mechanisms underlying sleep regulation remain elusive. We built machine learning (ML) models to predict genes based on their similarity to known sleep genes. Using a manually curated list of 109 labeled sleep genes, we trained a prediction model on thousands of published datasets, representing circadian, immune, sleep deprivation, and many other processes. Our predictions fit with prior knowledge of sleep regulation and also identify several key genes/pathways to pursue in follow-up studies. We tested one of our findings, the NF-κB pathway, and showed that its genetic alteration affects sleep duration in mice. Our study highlights the power of ML to integrate prior knowledge and genome-wide data to study genetic regulation of sleep and other complex behaviors.

## Introduction

Genetics impacts sleep. In humans, a handful of alleles are known to cause familial sleep disorders(1–8). However, most of these alleles are rare and have not been broadly implicated in sleep regulation in human populations. Genome-wide association studies (GWAS) identified more sleep-trait associated genes, but SNP-based heritability estimates are small. Few (if any) of these genes have been functionally validated(9–12). Many key features of sleep are conserved from invertebrates to vertebrates(13). Large-scale forward genetics screens in flies(14–16) and mice(17,18) have identified several genes whose alteration impacted sleep regulations. The two-process model(19,20) proposed that both circadian clocks and sleep homeostasis drive the sleep-wake cycle. Multiple studies have sought to identify key genes and proteins that regulate sleep homeostasis(21–25). Yet the molecular mechanisms underlying sleep regulation remain elusive.

Recent advances in ‘omics technology have led to increasingly large amounts of data generated each year. To date, the wealth of genome-wide datasets available have not been integrated to study the genetic regulation of complex physiology and behavior like sleep. Machine learning (ML) models have predictive power to classify samples based on hidden patterns in large datasets(26–29). Here, we applied ML to existing information with the goal of identifying genes and pathways involved in sleep regulation. Using a manually curated list of 109 labeled sleep genes, we trained a prediction model on thousands of published datasets, representing circadian, immune, sleep deprivation, and many other processes. Our model predicted 238 candidate sleep genes. Pathway enrichment analysis revealed the NF-κB pathway as a key factor in sleep regulation. We validated that activation of the NF-κB pathway in neurons indeed led to fragmented sleep in mice. In sum, we present an integrative *in silico* approach with the potential to identify genetic regulators of complex physiology and behavior.

## Results

### Defining sleep gene features

Our goal was to build a machine learning model to predict candidate sleep genes based on molecular features of known sleep genes. As a first step, we manually curated a list of known sleep genes (Table 1, hereafter referred to as ‘sleep genes’) through literature mining from PubMed and Scopus databases. Sleep genes were defined as genes reported to alter sleep traits, including sleep timing, sleep duration, and measurements of sleep quality from EEG in at least one animal model (flies or mammals).

**Table 1.**
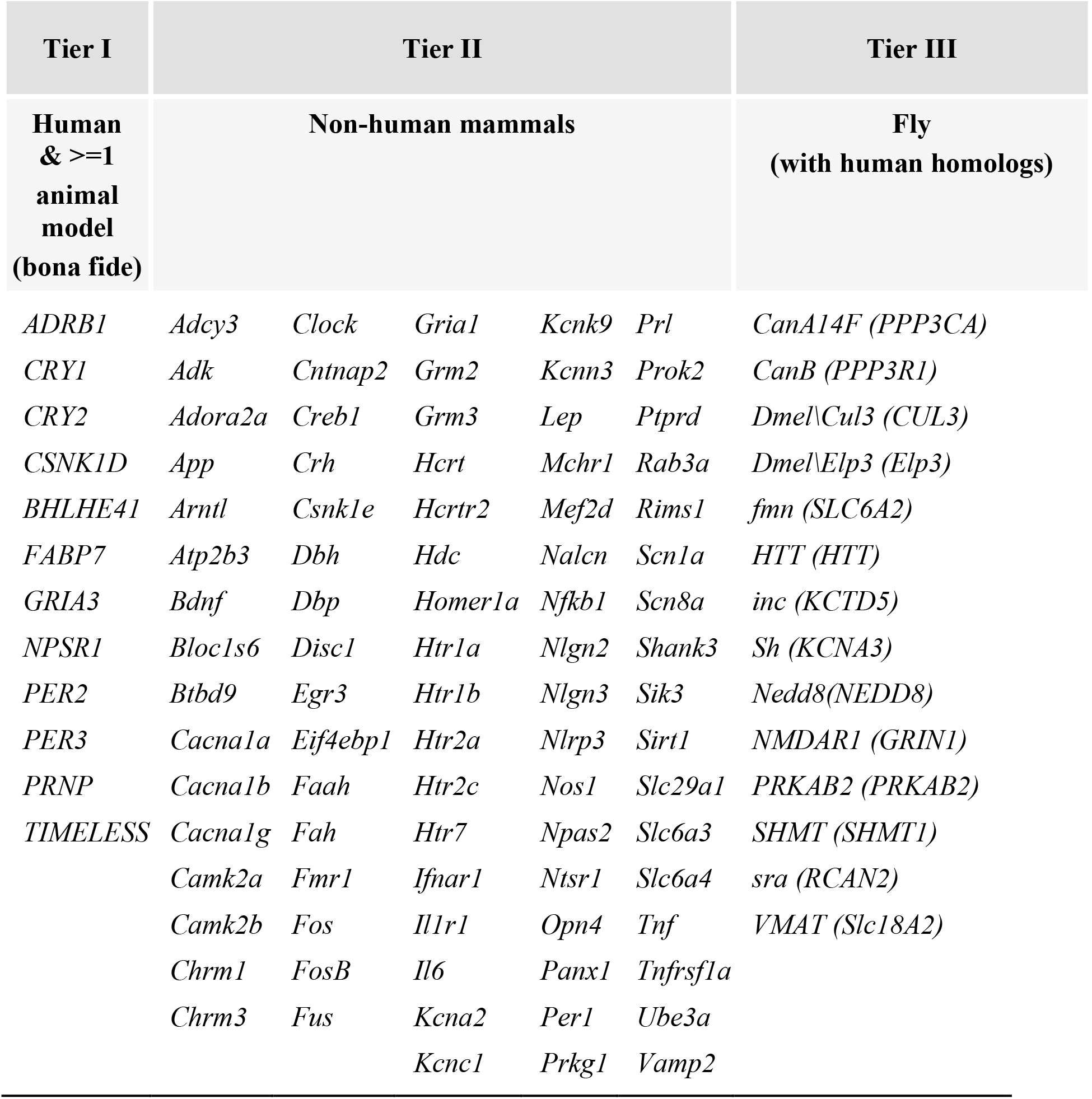
109 sleep genes curated through literature mining.

Next, we identified molecular features associated with these sleep genes. The lack of a strong molecular understanding of sleep regulation makes it difficult to know which types of information can be useful to predict sleep genes. To address this issue, we used two sources of information to define sleep gene-associated molecular features. The first source includes gene and protein knowledge from annotated gene set collections, including canonical pathways, gene ontology, transcription factor target genes, and protein-protein interactions. We applied the Jaccard index (JI), or the Jaccard similarity coefficient(30), to quantify the similarity of a gene to the exemplar sleep genes in the context of a given gene set collection (Fig 1A). Using the JI scoring method, we generated 19 features (S1 Table) representing the similarity of a gene to sleep genes in various molecular contexts.

**Fig 1.**
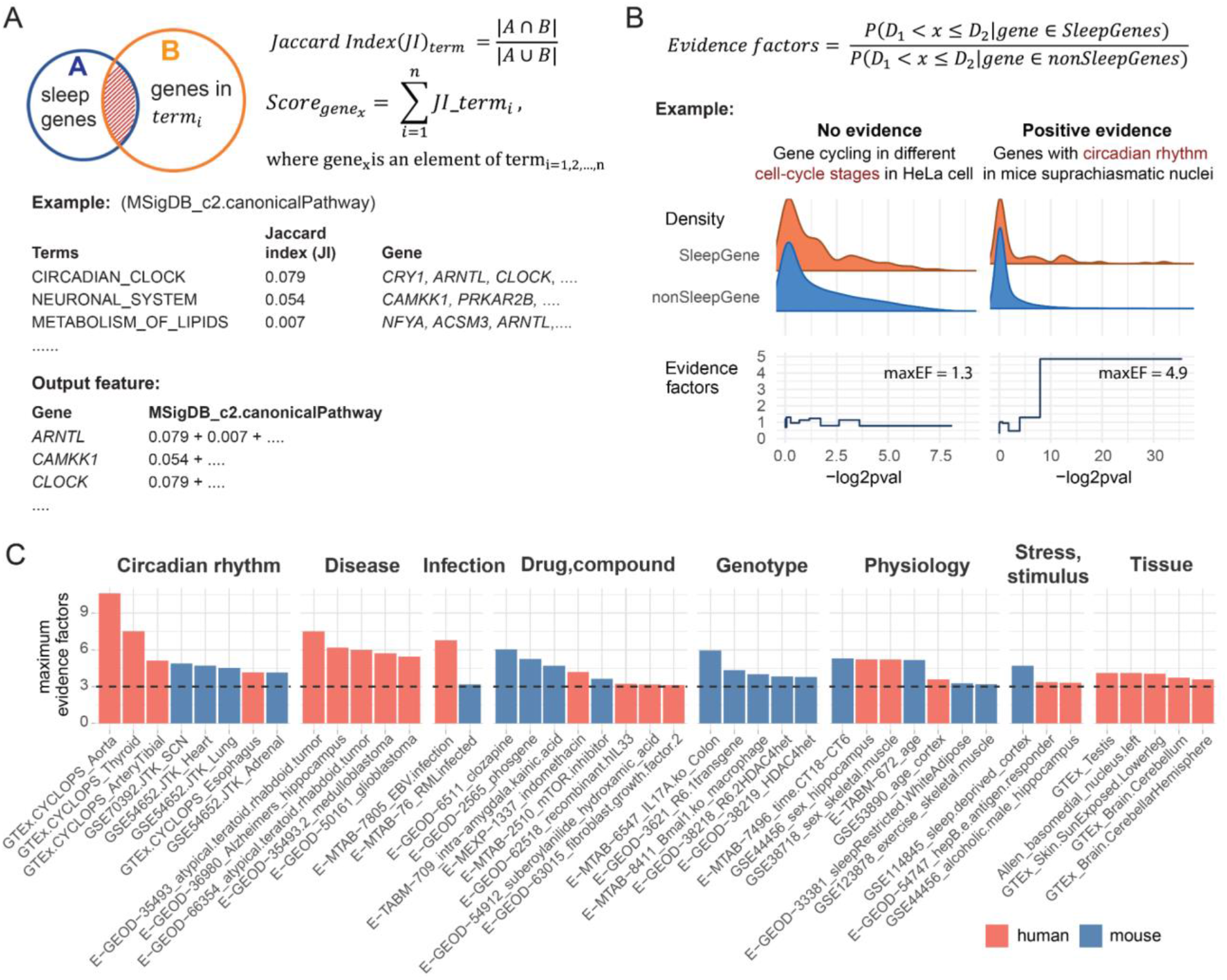
Defining sleep gene associated features. Two lines of information are used to define the sleep gene-associated molecular features. **A**. Gene and protein knowledge from annotated gene set collections. An example is shown here on how Jaccard index (JI) is used to score genes in a given gene sets collection. First, we calculate JI for all terms in the collection. Next, for each gene, we iterate through all terms. We added a term’s JI to the gene if this gene is an element of the term. The sum of JI represents the similarity index of a gene to sleep genes given this molecular context. **B**. Genome-wide datasets. Evidence factors are used to screen for datasets that show over-representation of sleep genes. Two time-series transcriptomics datasets are shown as negative and positive control for this screening method. We used −log2(p-value) as a significance score for rhythmic expression. Significance scores are split to form the sleep genes and non-sleep genes distribution. Evidence factors are calculated by comparing the proportion of genes in these two distributions within a bin. **C**. Genome-wide datasets enriched for sleep genes. Top 8 datasets with maxEF larger than 3 in each group are shown in the figure. Groupings include circadian rhythm, disease, infection, genotype, drug/compound, physiology, and stress/stimulus and anatomical specific transcripts/protein expressions. Y-axis of the bar plots show the maximum evidence factors for each dataset. Human and mouse samples are colored in red and blue, respectively.

The second source of information we used to define sleep gene-associated molecular features includes genome-wide profiling datasets. We used evidence factors(31,32) to identify genome-wide datasets most likely to be informative for the ML model. We evaluated 7,195 datasets for sleep-gene over-representation using maximum evidence factors (maxEF). In prior work, we applied evidence factors to identify a novel circadian transcriptional repressor in mice(33). We modified the application here to screen for datasets that show positive evidence for sleep genes (Fig 1B). To validate the concept, we tested two time-series datasets: (i) as a negative control, a transcriptomics profile of HeLa cells at different cell-cycle stages (GSE26922)(34) and (ii) a transcriptomics profile of mouse suprachiasmatic nucleus (SCN) across a 48h time-span (GSE70392) as a positive control for sleep gene regulation. These two datasets were selected as controls because the cell-cycle stage is not expected to be predictive of sleep, whereas circadian rhythms are intimately linked with sleep. The widely accepted two-process models of sleep regulation (19,20) and the alleles identified in families with extreme sleep traits(2,6,35) both support the roles of endogenous circadian rhythms in sleep regulation.

For the two control datasets, each gene was assigned a significance score for rhythmic expression using the published −log2(p-value). For each dataset, we built two distributions using this significance score. The 109 known sleep genes were used to form a sleep gene distribution. All remaining genes were used to form a non-sleep gene distribution. The evidence factors were computed by comparing the proportion of genes in these two distributions. If the two distributions are similar, maxEF is close to 1, which would indicate that there is no sleep gene over-representation in the dataset. In contrast, if the sleep gene distribution is different from the non-sleep gene distribution, maxEF would be much greater than 1. Evidence factors greater than 3 suggest positive evidence(31). Therefore, a cutoff of maxEF larger than 3 is set as an indicator of sleep gene over-representation in the dataset. As expected, we found no evidence of sleep gene over-representation in the cell-cycle time-series dataset (maxEF=1.3). Conversely, sleep genes were overrepresented in the mouse SCN time-series dataset (maxEF=4.9). Rhythmically expressed genes in this dataset were five times more likely to be sleep-versus non-sleep genes (Fig 1B), suggesting that circadian expression in the mouse SCN is a sleep-gene associated feature and should be incorporated in our ML model.

We screened through 7,195 genome-wide profiling datasets for positive evidence for sleep (maxEF>3) (S1 Fig & S1 Table). Datasets with the highest maxEF included circadian expression of genes in multiple tissues, and altered gene expression in several brain diseases. Sleep genes were also over-represented in datasets pertaining to Epstein-Barr virus infection, IL17A knockout in colon, clozapine treatment, sleep deprivation, and anatomically-specific datasets, including testis and human brains (Fig 1C & S2 Fig). In sum, we identified 72 datasets with high maxEF to provide computational and predictive efficiency for the ML model.

### Applying ML models to discover novel sleep genes

From the previous section, we selected 19 features from gene set collections using JI scoring methods and 72 features with high maxEF from genome-wide profiling datasets. Genes with >50% missing values from these 91 features were filtered out. One of the tier I sleep genes, *NPSR1*, was excluded as it had missing values in more than half of the selected datasets. With this information, we generated an input table with 17,841 samples (genes), including 108 labels (sleep genes), and 91 features for training the ML models.

ML models were built using Python packages scikit-learn(36) and Keras(37,38). We have curated 108 sleep genes (positive labels), but with no information or confidence on which genes do not regulate sleep (no negative labels). We applied a biased learning method to solve this problem of learning from <u>P</u>ositive and <u>U</u>nlabeled data(39,40). To do this, all non-labeled genes in the training set are treated as negative labels during the training process. In this case, the negative labels contained a mixture of true and false negatives, which led to weak, or low confidence, classifications for individual models. We subsampled the training sets with 7 training ratios (ranging from 0.2 to 0.8), each with 100 cycles, and made the final prediction results based on collections of all these weak-classifiers (Fig 2A).

**Fig 2.**
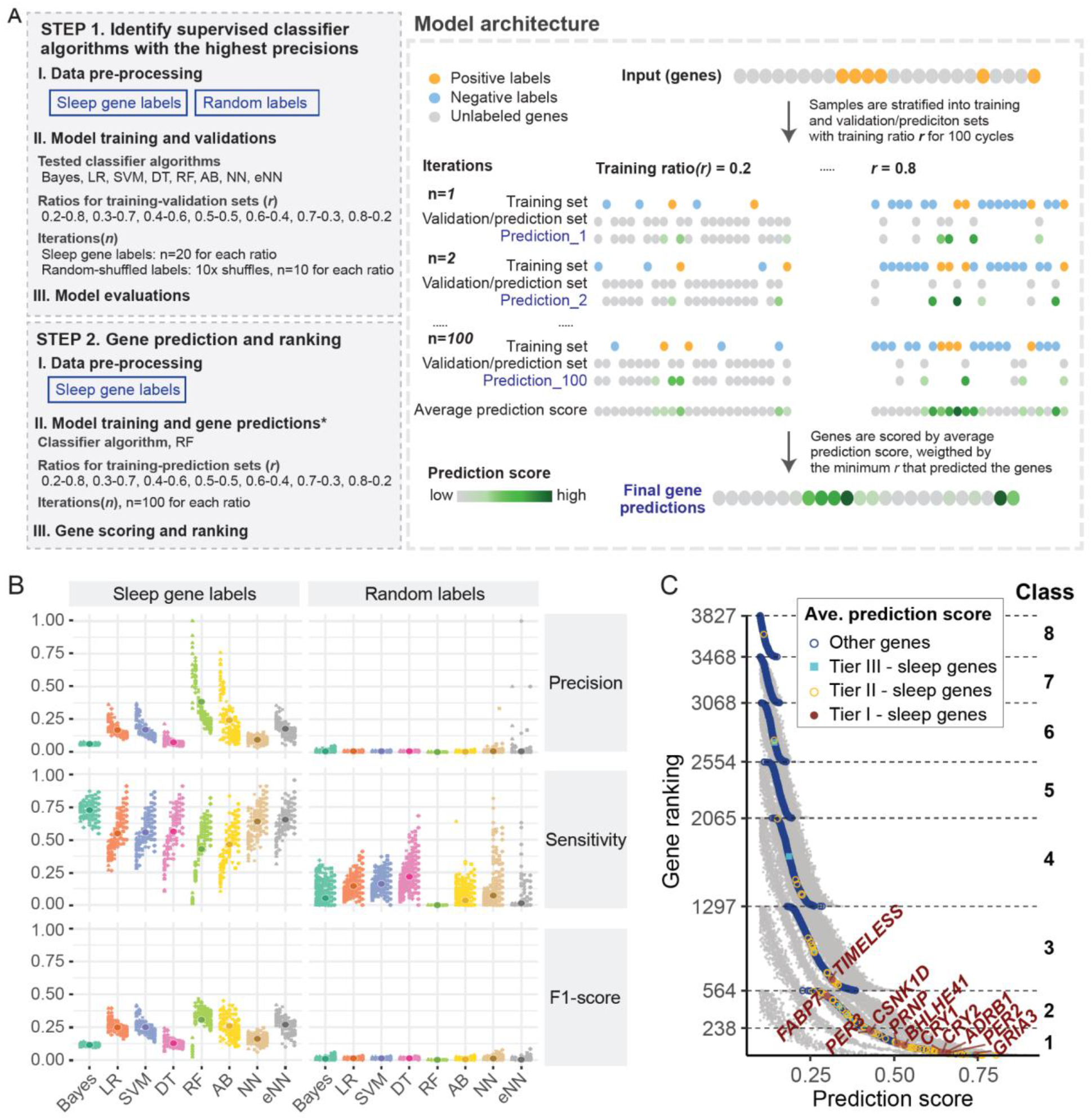
Building machine learning (ML) models to discover novel sleep genes. **A.** Workflows and model architecture for the ML models. We first tested the performance of 8 supervised classifier algorithms. Next, we selected algorithms with the best performance to build the sleep gene prediction models. Sleep gene prediction models are trained as shown in the model architecture. **B.** Precision, sensitivity and F1-score are used for model evaluations. The two columns indicate model performances using sleep gene labels and random-shuffled labels trained input. Each data point indicates performance from one iteration. Performances of different training ratios are shown in individual columns with different shapes, from left to right, ranging from 0.2 to 0.8. The average score for each classifier is shown as a circle at the center of each group. **C.** Prediction and ranking of sleep genes by random forest. Average prediction score of 0.1 is used as a cutoff for positive prediction. We classified all positive predicted sleep genes into 8 classes. Class 1 are the top 238 genes with average prediction scores larger than 0.413. Remaining predicted sleep genes are classified into class 2 to 8, based on the minimum training ratio that leads to positive predictions for the gene. For example, *TIMELESS* is classified into class 3 sleep genes as it is predicted when at least 30% genes are used as training sets. When smaller portions of genes are used in the training sets, less genes are predicted but with higher precision, and therefore with higher confidence. Points in grey show the prediction score from all training ratios. Average prediction scores for sleep genes validated in humans (Tier I), non-human mammals (Tier II) and drosophila (Tier III) are colored in brown, yellow and light blue, respectively. Remaining genes are colored in dark blue. Tier I sleep genes are labelled in the figure. *Abbreviations*. *Bayes*-naive Bayes, *LR*-logistic regression, *SVM*-linear support vector machines, *DT*-decision tree, *RF*-random forest, *AB*-adaptive boosting, *NN*-neural network, *eNN*-ensemble neural network.

There are numerous supervised classifier algorithms available, each with strengths and weaknesses. Our goal was to identify the highest-confidence candidate sleep genes for subsequent validation in animal models. We evaluated eight supervised classifiers seeking a model to maximize precision. False positives are much worse than false negatives, as validation experiments can take years. The tested classifier models included probabilistic (naive Bayes), linear regression (logistics and linear support vector machines), decision tree-based (decision tree, random forest and adaptive boosting), and neural networks (neural networks and ensemble neural networks). As part of our evaluation, we retrained all models with random-shuffled labels as inputs. This evaluation helped to minimize the chance that predictions are made based on random noise. Classifiers with high sensitivity in these random-shuffled label models were rejected.

Random forest and adaptive boosting performed best (AUC=0.97), followed by neural networks (AUC=0.96) (S3 Fig). Random forests had the highest precision, lowest sensitivity, and highest harmonic mean of sensitivity and precision (F1-score). Random forests also outperformed all other classifiers with regard to random-shuffled label models, with 0 for sensitivity, precision, and F1-score, suggesting high precision of the prediction results (Fig 2B). We therefore chose random forests as the classifier algorithm to predict sleep genes.

In total, 3,827 out of 17,841 genes were predicted as sleep genes (Fig 2C, S2 Table). We ranked these genes based on average prediction score, and separated them into 8 confidence levels based on the minimum training ratio that led to a positive prediction. We refer to the top 238 genes as class 1 candidate sleep genes. Of the class 1 candidate sleep genes, 63 were known sleep genes (Table 1) and 175 were novel predicted sleep genes. Sleep genes identified from human samples ranked higher in comparison to sleep genes identified from other mammals or flies, despite the fact that all sleep genes were weighted equally during the feature selection and model training steps. This suggested that our models’ predictions are able to detect human sleep genes and provide strong candidates for future study.

### Identifying pathways relevant to sleep regulation

The prediction model was intended to reveal molecular mechanisms or pathways that may be involved in the regulation of the sleep-wake cycle. We ran enrichment analysis (Reactome) with DAVID(41) using the 238 class 1 candidate sleep genes to explore the pathways enriched for sleep regulation. Pathways enriched by similar sets of genes are clustered into groups using Kappa similarity. We identified 19 enriched pathways (S4 Fig), 11 of them have at least 2 genes overlapped with the annotated genes from 4 GWAS pertaining to chronotype (9), overall sleep duration (12), insomnia (11) and daytime sleepiness (42)(Fig 3).

**Fig 3.**
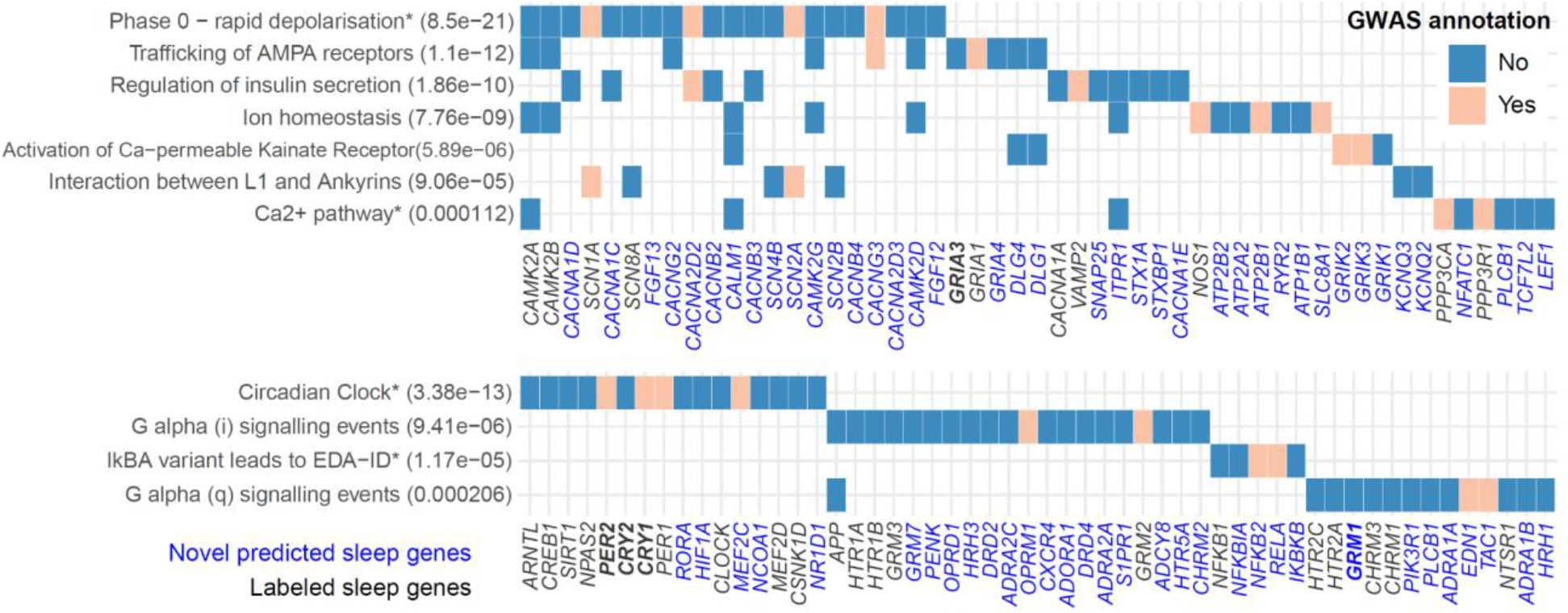
Enriched (Reactome) pathway from predicted sleep genes. Top enriched pathways with at least 2 genes overlap with sleep traits GWAS are shown in the figure. The full list of annotated pathways can be found in the S4 Fig. P-value for each annotated pathway is shown in the pathway labels. Pathways enriched by similar sets of genes are clustered into groups using Kappa similarity. For each cluster, pathway with the highest number of genes overlapped to GWAS (and lowest p-value if tied) is shown in the figure and marked with (*). Predicted genes overlapped to sleep traits GWAS are colored in beige and others are colored in blue. Sleep genes used to train the ML model are labeled in black and the predicted sleep genes are labeled in blue. Within each pathway, genes are ordered by their predicted rankings, from left to right. The five genes written in bold italic are the sleep genes that have been reported to regulate sleep in human studies.

Several of these pathways are neuron-related, including Phase 0 depolarization; ion homeostasis; Ca^2+^ pathway; trafficking of AMPA receptors and activation of Ca-permeable Kainate receptor. This is not surprising as neuronal involvement in rapid transition between sleep-wake states is well known(22,43–45). Our ML models proposed candidate sleep genes in each of these pathways that are yet to be explored. *CACNA2D2, SCN2A, CACNG3, ATP2B1, SLC8A1, GRIK2 and GRIK3* are supported by both ML models and GWAS data and represent attractive candidate genes for experimental validation.

Previous GWAS for sleep traits have reported enrichment of circadian rhythm and G-protein relevant pathways(9,12). Similar trends are observed here; circadian clocks and G alpha signaling events are among the top enriched pathways from our prediction models. Interestingly, two genes encoding opioid receptors, *OPRD1* and *OPRM1*, and a gene encoding the endogenous opioid peptides, *PENK*, are among the top candidate genes in G alpha(i) signaling events. Opioids are well known sedatives. Clinical studies have shown that a single dosage of opioid medication can significantly affect sleep architecture in healthy adults(46). Our prediction results suggest that among the three opioid receptors, the mu- and delta-receptors are more likely to play key roles in sleep regulation at the molecular level. This is in agreement with an *in vivo* study using opioid receptor agonists in feline models(47).

### Validation of a role for NF-κB activation in sleep regulation

We also found enrichment of pathways without prior association with sleep. In particular, a group of immune related genes, including *IKBKB*, *NFKB1*, *NFKB2*, *NFKBIA*, and *RELA*, were among the top enriched pathway clusters (Fig 3). These genes are key components of the proinflammatory NF-κB pathway in which *RELA* is a transcriptional factor and *IKBKB* is an upstream regulator of NF-κB activation. NF-κB transcription factors play critical roles in inflammation and immunity, as well as cell proliferation, differentiation, and survival(48). The direct and indirect triggers of NF-κB activation have been reported to cause circadian disruption(49). Sleep loss alters immune function and immune challenges alter sleep(50). Previous studies reported that *Nfkb1* (p50) knockout mice showed increased durations of slow-wave and rapid eye movement (REM) sleep(51). However, little is known about the direct effect of NF-κB activation on sleep homeostasis.

To test the effect of NF-κB activation on sleep, we used Camk2a^CreER^::R26-stop^FL^Ikk2^CA^ mice (Fig 4A, B) in which the stop^FL^ cassette prevents expression of the constitutively active *Ikbkb2* (Ikk2^CA^)(52) and the tamoxifen-inducible neural specific Camk2a^CreER^ recombinase(53) induces deletion of the stop^FL^ and thus expression of Ikk2^CA^(Fig 4C). IKK2 is a key component of the IKK complex which phosphorylates IkBα, leading to IkBα ubiquitination and proteasomal degradation(54). Upon degradation of IkBα, NF-κB is free to translocate to the nucleus, bind to DNA, and induce transcription of target genes. Therefore, Ikk2^CA^ expression leads to constitutive NF-κB activation, and these mice represent a genetic model of NF-κB pathway activation. We performed the PiezoSleep assay to assess the sleep-wake phenotypes.

**Fig 4.**
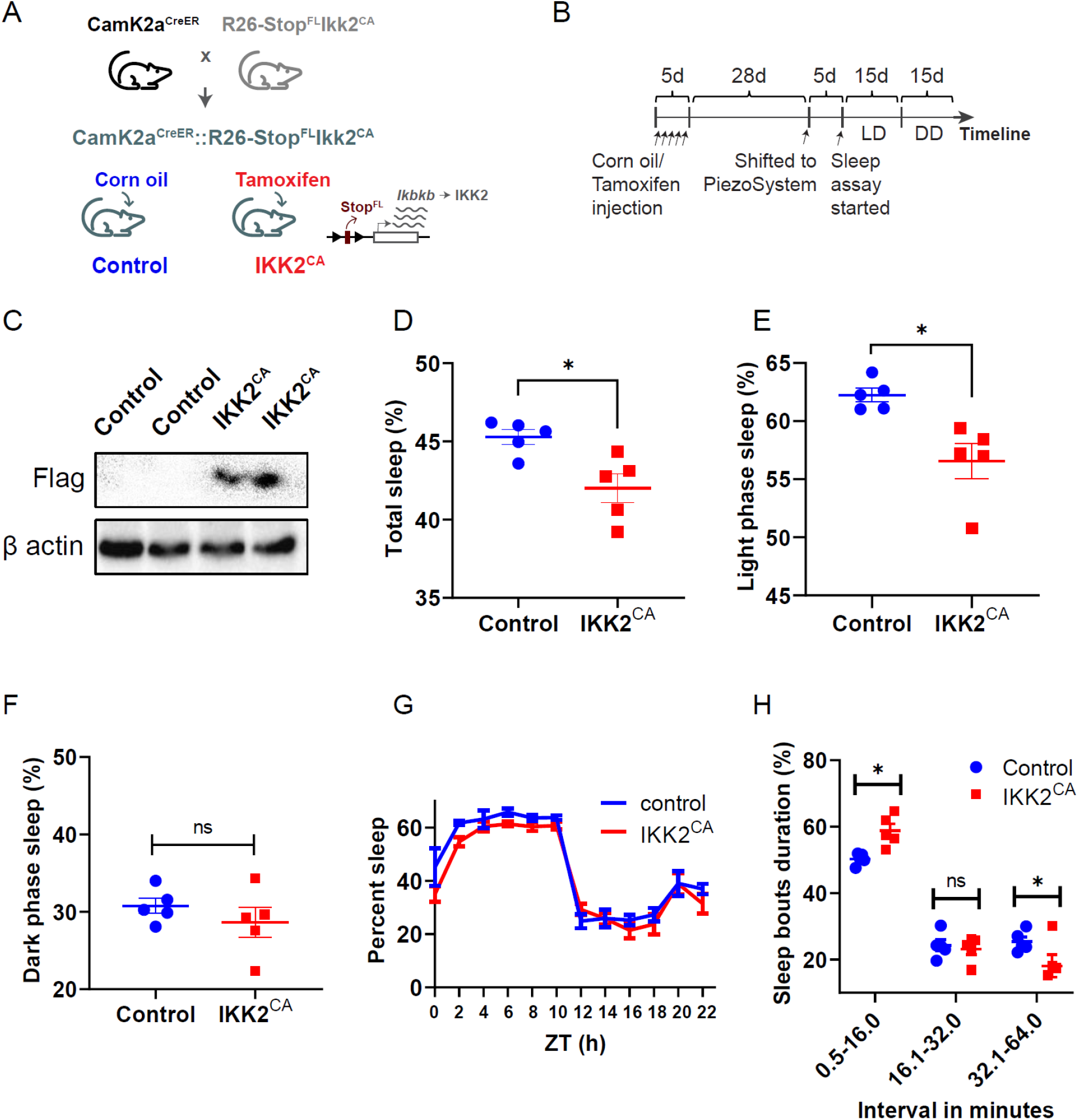
Sleep phenotyping in Ikk2^CA^ mice and control using piezoelectric sleep monitoring system. **A.** Camk2a^CreER^::R26-stop^FL^Ikk2^CA^ mice are used to test the effect of NF-κB activation on sleep. The stop^FL^ cassette in R26-stop^FL^Ikk2^CA^ mice prevents expression of the constitutively active *Ikbkb* (Ikk2^CA^) and the tamoxifen-inducible neural specific Camk2a^CreER^ recombinase induces neuronal specific deletion of the stop^FL^ and thus expression of Ikk2^CA^. **B.** Five doses of Tamoxifen and corn-oil are injected to Ikk2^CA^ and control mice, respectively, 4 weeks prior to the sleep phenotyping. Mice shifted to Piezoelectric sleep monitoring systems took 5 days for adaptation purposes. Sleep phenotyping is run for 15 days under LD (light on at 07:00-19:00, 250 lux), followed by another 15 days under DD. Five animals are used for each control and Ikk2^CA^ groups. **C.** Western blot data showed expression of flag-tagged Ikk2^CA^ in the Ikk2^CA^ mice but not control. Two out of five samples for each group are shown in this figure. **D-F.** The Ikk2^CA^ mice had a reduced (D) total sleep duration and (E) light phase sleep duration but no significant difference was observed in the (F) dark phase. **G.** Sleep reduction in the Ikk2^CA^ mice was observed during the light phase, from ZT0 to ZT10, in comparison to the control littermates. **H.** Ikk2^CA^ mice displayed more short bouts (0.5 to 16 min) (64.8 ± 0.57% in Ikk2^CA^ vs. 50.3±0.47% control) and less long bouts of sleep (≥ 32 min) (9.4 ± 3.4% in Ikk2^CA^ vs. 25.5 ± 2.1% in control) in comparison to controls. T-test is used for all statistical comparison. P-value < 0.05 is marked with (*). Abbreviation. *ns* - non statistical significance.

Compared to control mice, the Ikk2^CA^ mice had a reduced total and light phase sleep duration (Fig 4D&E, *t-test, p<0.05*). No significant difference was observed in the dark phase when mice are normally active (Fig 4F). The sleep reduction in the light phase in Ikk2^CA^ mice spanned from ZT0 to ZT10 (Fig 4G), when mice are typically inactive. Sleep bout duration has been used as indicators of sleep consolidation vs. fragmentation (55,56). Ikk2^CA^ mice displayed more short bouts and less long bouts of sleep, compared to controls (Fig 4H, *t-test*, p<0.05), indicative of sleep fragmentation. Taken together, when the NF-κB pathway is activated, mice exhibit sleep fragmentation especially during the inactive/resting phase.

## Discussion

Our computational approach predicted 238 genes and 11 biological pathways involved in sleep regulation. Predictions fit with prior knowledge of sleep regulation, and also identify several novel avenues to pursue in follow-up studies. We tested one of these, the NF-κB pathway, and showed that its genetic alteration affects sleep duration in mice.

A role for immunity in sleep regulation is known. Sleep changes in response to infection. Inflammatory mediators such as IL-1, TNF, and prostaglandins appear to have sleep regulatory properties(57). Our ML model suggested that the activation of NF-κB pathways, specifically through the phosphorylation of the IkBα complex, is a key regulator in sleep. We validated one of the predicted genes (*Ikbkb*) using a neuron-specific, constitutively activated IKK2 (Ikk2^CA^) mouse model. We found that Ikk2^CA^ mice have reduced sleep duration and shorter sleep bout duration than the controls. The decrease in bout length and reduced sleep duration during the inactivity phase suggests disruption in sleep consolidation and increased sleep fragmentation that may be relevant to human sleep. Sleep perturbations including fragmented sleep with frequent night-time awakenings and excessive daytime sleepiness are common in human patients with neurodegenerative diseases or cancer, and these daily disruptions are a major factor for sleep disorders(58,59).

Machine learning has been widely applied to integrate biological data in recent years. Multiple gene prioritization tools have been developed(60–62), but most are built on the hypothesis that causal variants or driver genes and pathways exist and thus may not be ideal for understanding genetic regulation in complex traits. We sought to identify candidate sleep genes that share similar molecular features to the known sleep genes. Key to this approach is the ability to define a comprehensive yet predictive set of features. Most ML models for gene prioritization draw from annotation resources (e.g., GO terms, MSigDB, GWAS catalog)(27,61), we applied a modified probabilistics method to screen and select sleep-relevant features from raw or processed genome-wide data. This allowed us to cull 73 features from thousands of datasets. We think this represents a general framework for integrating large amounts of genome-scale data to predict genetic regulators in other complex traits.

There are limitations to this study. The quality of the predictions from a ML model depends on the quality of the training labels (ie., sleep genes), the relevance of features to labels, and the amount of information available per sample (ie., gene). Missing information reduces sensitivity of the models. For example, the expression level of *NPSR1* is low or unmeasurable in most genome-wide studies. Therefore, it is not likely to be identified or recalled by the prediction model. Incorporating new information, whenever available, will improve performance. Our current screening methods are not sensitive for datasets with smaller numbers of genes/proteins (e.g., n < 1000) or with lower resolution (e.g., binary output), such as most proteomics or single cell studies. Alternative scoring or screening methods are needed to incorporate this information. Model evaluation results indicated that ensemble neural networks perform comparably to random forests, with higher sensitivity but lower precision. This suggested that neural networks may be useful to extend the candidate gene list, particularly for inferring sleep pathways.

In sum, we built ML models to predict clusters of genes based on molecular similarity to the known sleep genes. This approach successfully identifies key pathways that are involved in sleep regulation. In addition to our validation of the NF-κB pathway, a few of the top candidate genes, *Mef2c(63), GRM1(64),* and *Tac1(65)* were independently validated. Our study highlights the power of ML-based tools to integrate prior knowledge and genome-wide data to study genetic regulation of sleep and other complex behaviors.

## Material and Methods

### Data curations and preprocessing

Gene name conversion between species was done using the homologene function (v1.5.68) (homologeneData2 database updated on 2019 April)(66) integrated in the limma package (v3.40.6)(67). Gene aliases (both human and mouse) were converted to official gene symbols according to gene info downloaded from NCBI on 04/01/2020 (hereafter referred to as ‘gene_info_04012020’)(66). https://ftp.ncbi.nlm.nih.gov/gene/DATA/GENE_INFO/Mammalia/Homo_sapiens.gene_info.gz

#### Annotated gene set collections

GMT files, representings gene and protein knowledge from annotated gene set collections, were downloaded from MSigDB(68) (n=11) and Harmonizome(69) (n=7). Protein-protein interaction information was downloaded from BioGRID(70,71) (n=1), information in GMT format were extracted as follows - interaction types including colocalization, direct interaction, and association and physical associations, from human data, are used. Protein names are matched with the official gene name using gene_info_04012020.

Genome-wide profiling data were downloaded from different resources. In total, 7,195 data metrics are processed as described below.

#### I. Tissue-specific transcripts abundances (n=595)

Microarray data from human tissues are downloaded from BioGPS - GSE1133(72,73). Average values from each tissue are transformed with log2 to create data metrics (n=84). RNA-seq quantifications from human tissues are downloaded from GTEx(74). Average TPM from the same tissues are transformed with log2 to create tissue-specific transcript abundances data metrics (n=54). Additional RNA-seq data, transcript expression summarized at per gene(protein) level, are downloaded from Human Protein Atlas (HPA)(75). Log2 protein-transcripts per million (pTPM) are used to create data metrics (n=43). Brain region-specific transcripts quantifications (log2 transformed) from Allen Brain Map(76) are downloaded from Harmonizome(69). The mRNA expression data representing brain structures specific transcript abundances are used to create data metrics (n=414).

#### II. Tissue-specific protein abundances (n=30)

Mass spectrometry-based proteomics data from human adult and fetal tissue samples are downloaded from the Human Proteome Map(77). Normalized quantifications from the gene-level expression matrix are transformed with log2 to create data matrices (n=30).

#### III. Significance of circadian expressions (n=25)

Time-series data from mouse tissues are downloaded from GSE54652(78) and rhythmic signals are detected using Meta2D-JTK in MetaCycle(79). Transformed significant value, −log2(p-value), is used to create data metrics (n=12) that represent the significance of circadian expression in mouse tissues. Circadian expressions from human populations tissues, ordered by CYCLOPS(80), are downloaded(81,82). Transformed significance value, −log2(p-value) are used to create data metrics (n=13).

#### IV. Transcriptional profiles under perturbations or different physiological/pathological conditions (n=6,540)

15 datasets from Gene Expression Omnibus(GEO)(83) are downloaded and preprocessed manually. Absolute log2 fold-changes for each tested condition are used to create data metrics (n=46). In addition, 2,459 human and mouse processed datasets are downloaded from EBI expression atlas(84). Data metrics (n=6,494) are created using absolute log2 fold-changes for each tested condition.

#### V. Miscellaneous data (n=5)

Phosphorylation sites information is downloaded from qPhos(85). Number of tyrosine, serine/threonine, tyrosine and serine/threonine phosphorylation sites in each protein are used to create data metrics (n=3). Vertebrate homology information from 10 vertebrates, including human, chimpanzee, rhesus macaque, dog, cattle, rat, mouse, chicken, western clawed frog and zebrafish are downloaded from MGI(86). The number of vertebrates that share a homolog gene with humans is used to create a data metric (n=1) representing conservation of genes. Transcriptomics profiles from HeLa cells enriched for different phases of the cell cycle are downloaded from GSE26922(34). Rhythmic genes are detected using Meta2D-LS in MetaCycle(79). Transformed significant value, −log2(p-value), is used to create data metrics (n=1) that represent the significance of cell-cycle rhythmicity in the cell line.

### Preparing input for prediction models

#### SAMPLES

All human genes were used as samples for the model. All genes (61,527 unique genes) from gene_info_04012020 are used to create the human gene list.

#### LABELS

Labels (sleep genes) were manually curated through literature mining. The initial set of sleep genes was collected from a review paper(87). We then search for additional sleep genes using the keyword of ‘sleep’ in title, and ‘gene’ AND ‘model’ in the main text from PubMed and Scopus databases. A sleep gene was defined as a gene that has been reported to alter sleep traits in at least one animal model (flies or mammals) by genetic approaches. Altered sleep traits include changes in sleep timing (sleep phase), sleep duration and other measurements of sleep quality from EEG (e.g. slow wave activity, NREM/REM ratio, number of sleep bouts and sleep latency). We divided the list into 3 tiers. Tier I includes “bona fide” sleep genes that harbored a causal mutation in any human sleep traits and were validated in animal models. Their roles are conserved across species. Tier II genes have evidence from any non-human mammalian model system. Tier III genes were discovered to change sleep traits in *Drosophila* but not in vertebrates yet. Given that only a limited number of tier I genes were characterized, tier II and III were included to build the model with the same weight as tier I genes. Sleep genes are updated till 8/13/2020 for this analysis.

#### FEATURES

Sleep gene associated molecular features were built using two lines of information.

##### I. Gene set collections

A total of 19 gene set collections are downloaded and prepared as mentioned above. One feature is created from each gene set collection. Genes are scored by their similarities to the list of curated sleep genes under the biological context, for example, similarities of the shared molecular pathways between a gene with the curated sleep genes. To do this, we first calculated the Jaccard Index (JI), or the Jaccard similarity coefficient, for each term in a given gene set collection. Assuming that the curated sleep genes are set A and the genes assigned to the term (e.g. circadian clock) are set B, JI is calculated by dividing the number of overlapped genes between A and B to the number of all unique genes in A and B. Next, for all genes in set B, we updated the gene score by adding in this JI value to the gene score. These two steps are repeated for every term in a given gene set collection. By the end of these calculations, we obtained a numeric vector with the sum of JI score for each gene. This created a feature representing the overall similarities of a gene with the labeled sleep genes, under the biological context of the gene set collection information.

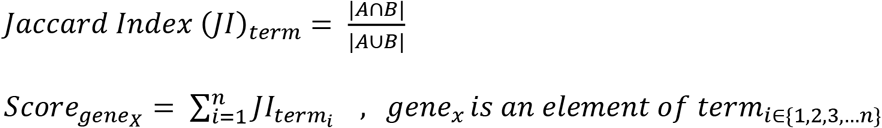

##### II. Genome-wide profiling datasets

Genome-wide profiling datasets are collected and pre-processed as mentioned in the data curations and preprocessing section. Evidence factors were used to evaluate the degree of sleep genes over-representation in these processed data metrics. We first split the samples (genes), using the sleep genes to form a sleep gene distribution, and the remaining genes to form the non-sleep gene distribution. Evidence factors are calculated by comparing the proportion of genes in these two distributions, within a bin (between D_1_ and D_2_). As the distribution of genes is sparse, there are chances to have empty bin(s) if we fix bin size by value. To solve this problem, we binned the samples(genes) by setting a minimum percentage of genes in each bin. To do this, the data metrics are first split by 100 equal breakpoints. We then repeatedly merge the neighboring bins until a bin has at least 10% sleep genes and at least 1% non-sleep genes, or at least 10% non-sleep genes and at least 1% sleep genes. For each (merged) bin, we calculated evidence factors by dividing the proportion of genes in sleep genes to the proportion of genes in non sleep genes. We used maximum evidence factors (maxEF) from a dataset as an index to select sleep gene relevant data. If the two distributions are similar, we will have maxEF near to 1, indicating no sleep gene over-representation in the data metric. In contrast, if the sleep gene distribution is different from the non-sleep gene distribution (e.g. skewed right tail in the sleep gene distribution, highly enriched for certain range of values), we will have higher maxEF, suggesting sleep gene over-representation in the data metric. For each data metric, we set a cutoff of at least 25% labels that must present with a real value to ensure sufficient sleep genes used to form the sleep gene distribution; and the range of the data metrics must have more than 3 steps to ensure sufficient resolution. Else, we skipped the maxEF calculation for this data metric and showed the maxEF value with ‘NA’. Data metrics that show positive evidence (maxEF>=3) are selected and used as features to train the prediction models. Most machine learning algorithms assume that all features are independent. To remove features that are highly correlated, we ran pairwise correlation coefficients of all data pairs. If two data metrics have correlation coefficient higher than 0.8, the data metric with lower evidence factors is excluded.

### Building machine learning models to predict sleep genes

#### INPUT

Samples and features are curated as mentioned above. We then filtered out samples (genes) with >50% missing values. One of the labels (sleep gene), *NPSR1*, was removed. In summary, an input table with 17,841 rows (genes) and 91 columns (gene-associated features), with 108 labels (sleep gene) was used to build the prediction models.

#### DATA PREPROCESSING

Data preprocessing is done using Python - sklearn.impute and sklearn.preprocessing package. Missing values from the input are imputed with mean value and rescaled with standard score (z-score). Curated labels (sleep genes) are replaced by ‘1’ and the remaining samples(genes) are replaced by ‘0’.

#### MODEL ARCHITECTURE

We have curated 108 sleep genes (positive labels), but with no information or confidence on genes that do not regulate sleep (no negative labels). This raised the problem of learning from <u>P</u>ositive and <u>U</u>nlabeled data (PU learning). We applied a biased learning method(39,40) to solve this problem. To do this, all non-labeled genes are treated as negative labels during the training process, and prediction results are made based on ensembles of numerous of these weak-classifiers. In other words, we first subsampled our samples (genes) into a smaller subset, with the same proportion of positive and unlabeled samples in the training and prediction sets. Samples in the training set are used to train the prediction models, sleep genes are marked as positive labels and all other genes are marked as negative labels (all other genes in the subset). In this case, the negatively labelled samples are expected to contain a mixture of true or false negative labels, hence, resulting in weak-classifiers. We repeated this process for 100 times and made the final predictions based on average performance from all cycles.

We have only a small number of labels (sleep genes, n=108) in comparison to other samples (non-labeled genes, n=17,733). Yet, these labels are not found completely at random. In other words, most sleep genes are identified based on our existing knowledge of sleep regulations; therefore, these genes are not distributed randomly (or equally) in all sleep-relevant pathways. As an example, 13 out of 108 sleep genes are parts of the circadian clock pathway. To increase the randomness of subsampling labels, as well as maintaining the best performance (sensitivity and precision), we trained the ML model with different proportions of training input, ranging from 0.2 to 0.8. By doing this, we increase the combinations of samples used in the training and prediction sets, and therefore expected to have more robust predictions.

#### MODEL EVALUATION

Eight supervised classifier algorithms were built to find the best supervised classifier that fits our prediction. The evaluated classifiers included probabilistics models (naive Bayes), linear regressions models (logistics and linear SVM), decision trees (decision tree, random forest and adaptive boosting) and neural networks (neural networks and ensemble neural networks). All machine learning models, except neural network and ensemble neural network, were built using scikit-learn (v0.22.2)(36). Default parameters were used, except mentioned below. Logistics regression (max_iter=1000), decision tree (max_leaf_nodes=12), random forest (max_leaf_nodes=12), adaptive boosting (max_leaf_nodes=4, algorithm=”SAMME”, n_estimator=200). Neural networks and ensemble neural networks were built using Tensorflow (v2.2.0)(38) and Keras (v.2.4.3)(37). Neural networks were built using sequential models with two hidden layers (12 and 6 nodes each, both activated by ‘relu’ function). Final outputs were activated by the ‘sigmoid’ function. Ensemble neural network had the same setting as the neural network, except we ran the model for 20 times for each subsampling input, and made the final predictions by major voting (>50%, or more than 10 times). Input to train prediction models are prepared as mentioned above. For naive Bayes classifiers, we transformed the input with principal component analysis to ensure the conditional independence between features. The ratio of labeled and unlabeled samples is skewed. To balance the weights of positive and negative labels to roughly 1:1, we assumed the total number of genes is around 20,000. The weight is calculated by using half of the total number of genes (10,000) divided by the number of labels. Samples (genes) are randomly split into training and testing sets, stratified by the same percentages of labels in the training and testing sets. This process is repeated for 100 cycles for each of the 7 training ratios, where training ratios include 0.2, 0.3, 0.4, 0.5, 0.6, 0.7 and 0.8. For each iteration, samples assigned to the training set are used to train the models. Samples in the testing set fit into the trained models, and the classes predicted by the “predict_classes” function are used to calculate the confusion matrix. Models are evaluated with precision, sensitivity and F1-score (harmonic mean of precision and sensitivity). Raw prediction scores, or the probability of a gene predicted as sleep gene by the ML model, calculated by the “predict_proba” function were recorded for genes assigned to the prediction set (not including linear SVM, as the raw prediction scores from linear SVM were discordant with the binary predictions). The average prediction score from all iterations was calculated and used to plot the sensitivity-precision plots.

#### RANDOM LABELS

To avoid a model that makes predictions based on random noise, we removed all existing labels and randomly assigned the same number of labels to the remaining samples (only samples that were not originally labelled). As features are built based on sleep genes labels but not the random labels, the randomly assigned labels are not likely to be recalled using these sets of features, unless called by random noise. Therefore, we selected models that have low sensitivity with the random labels input.

#### PREDICTION MODEL

The prediction model is built with random forest classifiers using scikit-learn (v0.22.2)(36). Default parameters were used, except max_leaf_nodes is set to 12. As described in the previous section, the weights of the labels are calculated using 10,000 divided by the number of labels. Samples (genes) are randomly split into training and testing sets, stratified by the same percentages of labels in both the training and testing sets. This process is repeated for 100 cycles for each of the 7 training ratios, where training ratios include 0.2, 0.3, 0.4, 0.5, 0.6, 0.7 and 0.8. For each iteration, samples assigned to the training set are used to train the models. Labels from the prediction set are removed; all samples in the prediction set are fit into the trained models for prediction. Raw prediction score is recorded for genes assigned to the prediction set. Prediction score less than 0.1 is set to 0 to reduce noise. The average prediction score from all iterations was calculated. We observed a linear correlation between training ratios with sensitivity, and an inverse correlation with precision (Fig2B). In fact, the smaller the training samples are, the fewer genes are predicted as sleep genes, but a larger portion of these predicted genes are indeed sleep genes (higher precision). For this reason, we weighted the final ranking of candidate sleep genes with the minimum training ratio that leads to a positive prediction (min(*r*)). A gene predicted as sleep genes in models trained by only 20% of the samples (genes) will rank higher than a gene only predicted as sleep genes in models trained by 80% of the samples. The final prediction score is calculated as:

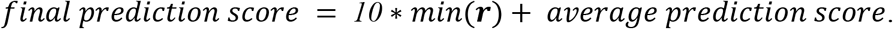

Genes with consistently high prediction scores in all training ratios (average prediction score >0.413, aka the top 238 genes) are grouped as the class I predicted genes.

### Exploring sleep traits GWAS data

Summary statistics from 4 self-reported UK Biobank sleep traits GWAS (n=453,379), including chronotype(9), overall sleep durations(12), daytime sleepiness(42) and insomnia(11), were downloaded from Sleep Disorder Knowledge Portal (SDKP)(88). FUMA’s SNP2GENE process(89) is used to run gene annotations. SNPs are mapped to genes using the posMap, eqtlMap and ciMap methods, with default parameters. Mapped GWAS genes overlapped with the top predicted sleep genes are marked in Table S2.

### Pathway enrichment analysis

The 238 class 1 candidate sleep genes are used for pathway enrichment analyses using DAVID (41) - Reactome pathway database. Pathways are then clustered using kappa similarity in DAVID (kappa similarity threshold>0.5). We filtered out pathways with less than 5 genes or Bonferroni adjusted p-value larger than 0.1.

### Mice

The R26-stop^FL^Ikk2^CA^ transgenic mice (Stock No: 008242) and Camk2a^CreER^ transgenic mice (Stock No: 012362) were both obtained from The Jackson Laboratory. The Camk2a^CreER^ and R26-stop^FL^Ikk2^CA^ mice were crossed and housed under 12:12-h light: dark (LD) cycle within the University of Florida communicore facility and fed and watered *ad libitum*. Animal care and experimental procedures were approved by the Institutional Animal Care and Use Committee at University of Florida following the Guide for Care and Use of Laboratory Animals of the National Institute of Health (IACUC# 202110057).

### Tamoxifen Injection

For tamoxifen inducible Ikk2^ca^ knock-in activation, Camk2a^CreER^::R26-Stop^FL^Ikk2^ca^ transgenic mice were generated by crossing Camk2a^CreER^ mice with R26-stop^FL^Ikk2^CA^ mice. Tamoxifen (TAM) (#T5648; Sigma-Aldrich, St. Louis, MO) was dissolved in corn oil (#C8267, Sigma-Aldrich, St. Louis, MO) at a concentration of 20mg/mL. 10-12 weeks-old male mice (n=5 for each group) were dosed at 75 mg/kg body weight (TAM or corn oil) intraperitoneally once every 24 h for a total of five consecutive days. The sleep assay began 4-weeks after tamoxifen injections when constitutively active IKK2 expression was induced in this model.

### Western blot

Brain tissue lysate preparation and immunoblotting analysis were performed using anti-Flag (65, Sigma-Aldrich, St. Louis, MO) antibody. Briefly, brain tissue was snapped frozen and lysed in the RIPA lysis buffer containing cocktails of proteases inhibitors (Roche) and phosphatase inhibitors (Sigma). Western blot was performed to determine Flag-tagged Ikk2^CA^ activation.

### Sleep assay

The piezoelectric sleep monitoring system (PiezoSleep version 2.11, Signal Solutions, Lexington, KY), is a highly sensitive, non-invasive, high throughput and automated piezoelectric system, which detects breathing and gross body movements to characterize sleep patterns in unsupervised sleep/wake recordings(90,91).

For each experiment, 5 tamoxifen or 5 corn-oil injected Camk2a^CreER^::R26-stop^FL^Ikk2^CA^ mice were individually housed in PiezoSleep cages with a sensor inside a temperature, humidity, and light controlled box. The first 3-5 days of recording was considered as the acclimation period to the piezo device. The 12h light/12h dark (LD) cycle (light on at 07:00 to 19:00; 250 lux) was performed for the 15-days LD followed by the next 15-days of 12h dark/12h dark (DD) with *ad libitum* access to food, water and nesting material. Sleep data were analyzed for multiple sleep traits of individual mice using sleepstats2p18 (Signal Solutions, Lexington, KY).

## Acknowledgements

We thank Andrew I. Su and Casey S. Greene for technical assistance; Ying-Hui Fu, Kottyan C. Leah, Christian I. Hong, Tongli Zhang, members of Hogenesch lab and Smith lab for thoughtful discussions.

## Author Contributions

Y.Y.L., G.W., M.D.R., L.J.F., R.C.A., D.F.S., A.C.L., J.B.H. designed the research; Y.Y.L., G.W., L.J.F., N.Y.C. collected and pre-processed data; Y.Y.L., G.W., M.D.R., R.C.A., J.B.H. contributed the analytic tools; M.E. and A.R.M. performed experiments and interpret data; Y.Y.L., M.E., G.W., M.D.R., L.J.F., A.C.L., J.B.H. wrote the manuscript.

## Data and materials availability

Gene set collections is available at MSigDB http://www.gsea-msigdb.org/gsea/msigdb/index.jsp, harmonizome https://maayanlab.cloud/Harmonizome/, protein-protein interactions information is available at BioGRID https://downloads.thebiogrid.org/; tissue specific expression data is available at BioGPS http://biogps.org/downloads/, GTEx https://gtexportal.org/home/datasets, HPA https://www.proteinatlas.org/about/download, Allen Brain Map https://human.brain-map.org/; protein abundances measurements is available at http://www.humanproteomemap.org/index.php; circadian expression is available at http://circadb.hogeneschlab.org/; transcriptomics profile is available at GEO https://www.ncbi.nlm.nih.gov/geo/, EBI ArrayExpress https://www.ebi.ac.uk/arrayexpress/; phosphorylation sites information is available at http://qphos.cancerbio.info/; statistical summary from sleep traits GWAS is available at https://sleep.hugeamp.org/; GWAS gene annotation is run using webtools FUMA https://fuma.ctglab.nl/; pathway enrichment analysis is run using https://david.ncifcrf.gov/. Code to prepare input features and run the machine learning predictions can be found at https://github.com/yyenglee/ml-sleep.

---

**S1 Fig.**
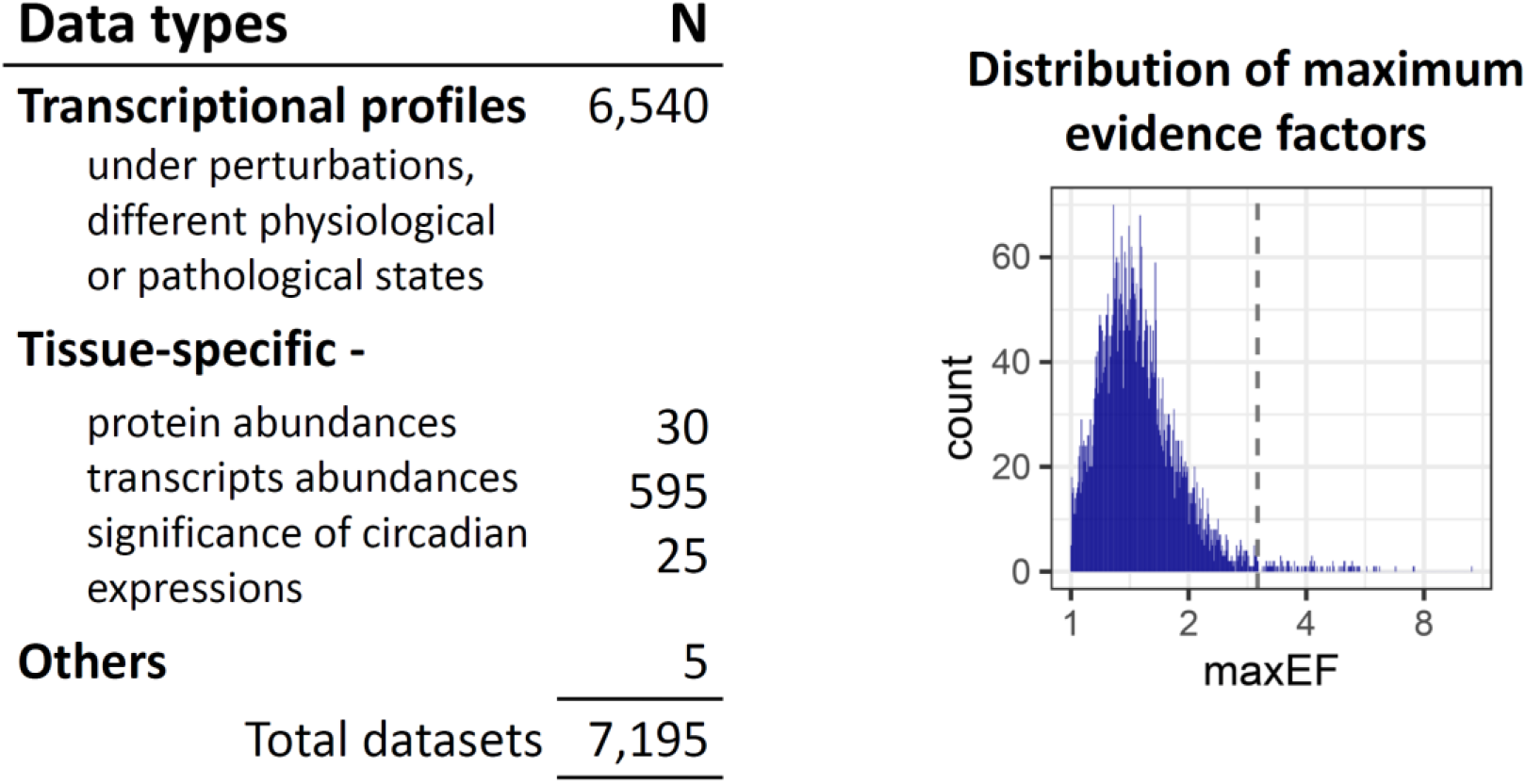
Summary of the genome-wide profiling datasets tested with evidence factors. Table above showed the number of data metrics tested for over-representation of sleep genes using evidence factors. The histogram showed the distribution of maximum evidence factors of all tested data metrics. A cutoff of 3 (represent positive evidence) is used to select features to train the sleep genes prediction model.

**S2 Fig.**
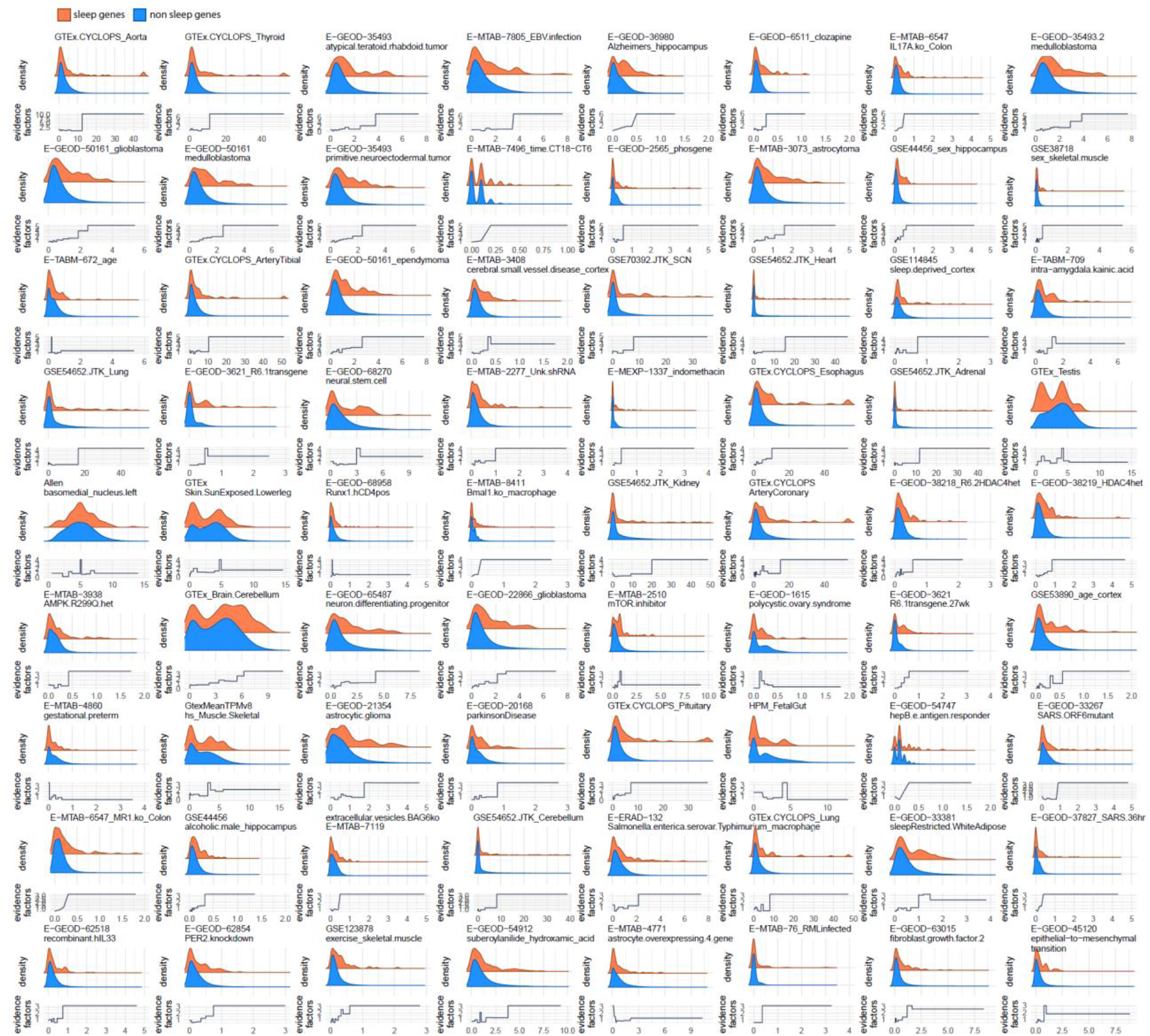
Evidence factors for the 72 selected genome-wide data metrics. Evidence factors are used to screen for genome-wide datasets enriched for sleep genes. In this figure, a data metric is represented by two density plots (top) and a line plot (bottom). The orange and blue distributions are generated using sleep genes and non sleep genes, respectively. The X-axis of the plot represents the range of value for this data metric, Y-axis of the line plot shows the evidence factors.

**S3 Fig.**
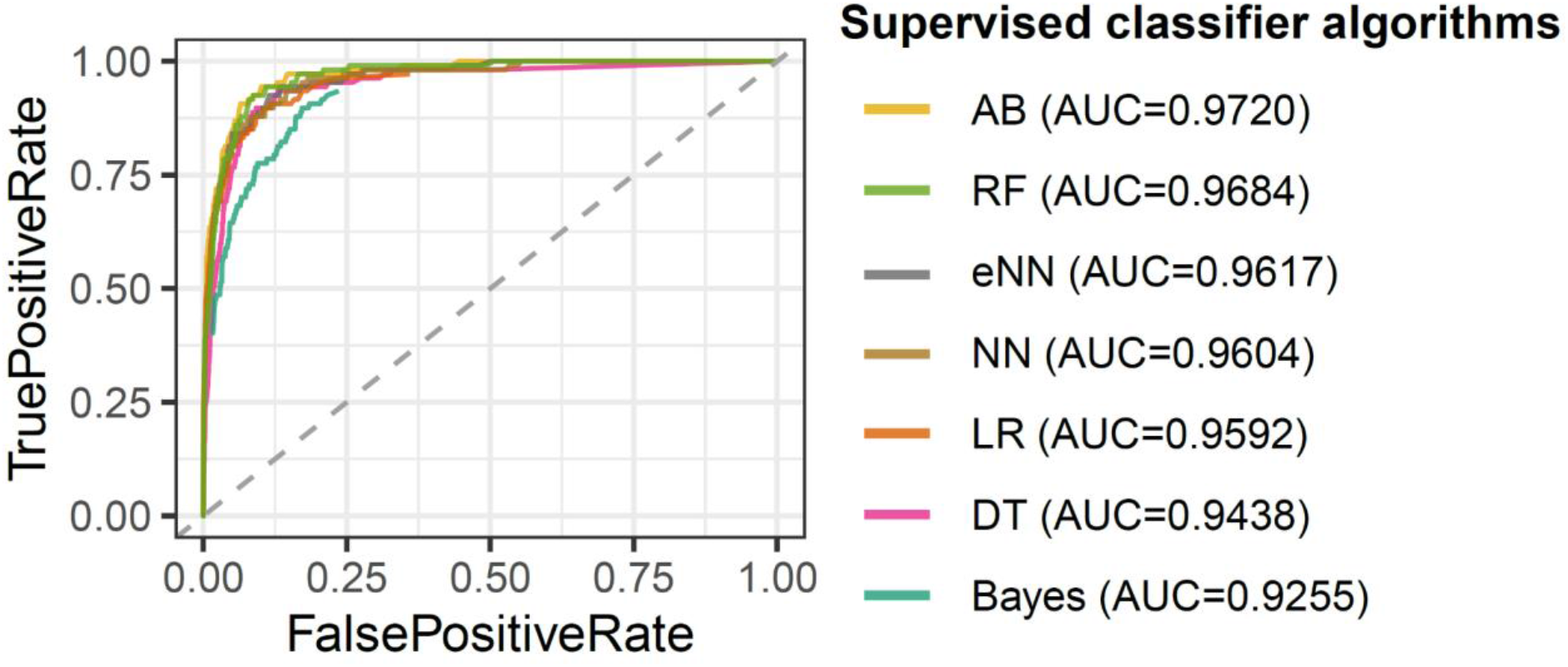
Receiver operating characteristic curve for tested classifier. Receiver operating characteristic (ROC) curves for each tested classifier (except linear SVM, as the probabilities scores are reported to be different from the classification predictions) are shown in the figure. Areas under curve (AUC) for each classifier are listed in the legends. *Abbreviations*. *Bayes*-naive Bayes, *LR*-logistic regression, *SVM*-linear support vector machines, *DT*-decision tree, *RF*-random forest, *AB*-adaptive boosting, *NN*-neural network, *eNN*-ensemble neural network.

**S4 Fig.**
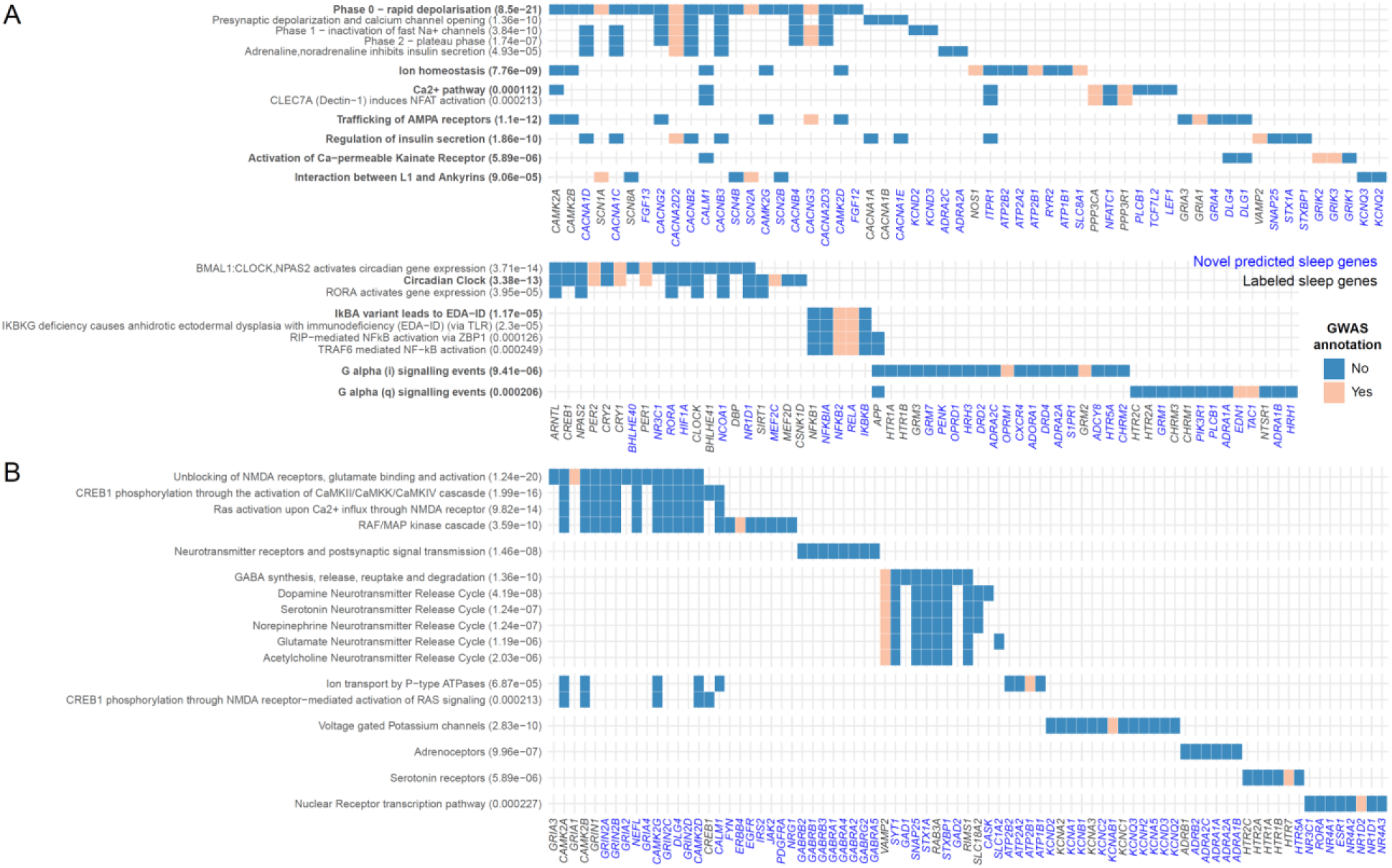
All enriched (Reactome) pathways from the top 238 predicted sleep genes. A. Pathways with at least two genes overlap with sleep traits GWAS mapped genes. B. Pathways with one or no genes overlap with sleep traits GWAS mapped genes. Pathways annotated by similar sets of genes are clustered with Kappa similarity threshold >0.5. Pathways in the same cluster are ordered by p-value, from top to bottom. Genes overlapped to sleep trait GWAS are colored in beige, others are colored in blue. Gene names labeled in black are sleep genes used as labels to train the ML model; gene names labeled in blue are the novel candidate sleep genes. Within each pathway, genes are ordered by their prediction rankings, from left to right.

## Supporting Information

**S1 Data. Known sleep gene curated through literature mining.**

**S1 Table. Curated datasets used to build ML models.**

**S2 Table. Ranking of sleep genes by random forest, with GWAS annotated genes annotation.**

