## Supplementary material for "Machine learning approaches to identify sleep genes": S1 Data

**Supplementary Data 1 - Known sleep gene curated through literature mining**

| **Tier** | **HGNC** | **References** | **Models** | **Genotype** | **Phenotype descriptions** |
| --- | --- | --- | --- | --- | --- |
| I | *ADRB1* | (Shi et al. 2019) | human, mice | hADRB1-A187V | Shorten REM and nonREM sleep;  Shorter sleep bout but not episode duration. |
| I | *BHLHE41* | (He et al. 2009) | mice | hDEC2-P385R | Short sleep. |
| I | *CRY1* | (Patke et al. 2017) | human patient/human dermal fibroblast/mice CRY1/2 deficient fibroblast | hCRY1-c1657+3A>C (5' splice site of exon 11) | Sleep phase delayed. |
| I | *CRY2* | (Hirano et al. 2016) | mice | hCRY2-hA260T | Familial advanced sleep phase (FASP);  Shortened circadian period. |
| I | *CSNK1D* | (Xu et al. 2005) | mice, drosophila | hCK1δ-T44A | Mice: advanced sleep phase  Drosophila: delayed sleep phase |
| I | *FABP7* | (Gerstner et al. 2017) | human, mice, drosophila | hFABP7-T61M, rs2279381 drosophila, mice  knockout(mice) | Shorter total sleep time and sleep bout duration;  Increased sleep bout frequency, this phenotype is significant only during the activity phase for knockout mice. |
| I | *GRIA3* | (Davies et al. 2017) | mice | hGRIA3-A653T | Fewer short bouts of sleep than the wild-types. |
| * | *GRM1* | (Shi et al. 2020) | human, mice | hGRM1-R889W  hGRM1-S458A | Familial natural short sleep.  Mice: shorter sleep duration. |
| I | *NPSR1* | (Xing et al. 2019) | human, mice | hNPSR1-Y206H | Shorter sleep duration. |
| I | *PER2* | (Vanselow et al. 2006) | fibroblast | hS662G | Short period/advanced phase. |
| I | *PER3* | (L. Zhang et al. 2016) | mice | hPER3-P415A/H417R transgenic mice | Advanced sleep phase. |
| I | *PRNP* | (Medori et al. 1992; Tobler et al. 1996) | human | hPRNP-D178N | Familial fatal insomnia. |
| I | *TIMELESS* | (Kurien et al. 2019) | mice | CRISPR-edited R1081X in mPER | Advanced phase of sleep-wake behavior;  Altered phase responsiveness to timed light pulses. |
| II | *BTBD9* | (DeAndrade et al. 2012) | mice, drosophila | B-geo gene trap vector at sixth intron (5th in human) | Sensory alterations likely to be limited to the rest phase;  Decreased sleep and increased wake times during the rest phase;  Decreased slow-wave sleep. |
| II | *DISC1* | (Jaaro-Peled et al. 2016) | mice, drosophila | expressing human DISC1 | Longer duration of wakefulness;  NREM sleep & REM sleep shorter;  No change in total activity level. |
| II | *ADCY3* | (X. Chen et al. 2016) | mice | knockout | Increase percentage of REM sleep. |
| II | *ADK* | (Palchykova et al. 2010) | mice | ADK transgene encoding the cytoplasmic isoform of ASK in the brain,  but lack the nuclear isoform of the enzyme | Awake 59 min more per day;  Spent significantly less time in REM sleep. |
| II | *ADORA2A* | (Huang et al. 2005) | mice | knockout | Normal sleep phase and duration;  Loss increases in waking after caffeine. |
| II | *APP* | (Huitrón-Reséndiz et al. 2002) | mice | hV711F, result in overexpression hβAPP | ~25% less REM sleep/24hr. |
| II | *ARNTL* | (Ehlen et al. 2017) | mice | knockout | Increase in total sleep time;  Sleep fragmentation and EEG delta power attenuated compensatory response to acute sleep deprivation. |
| II | *ATP2B3* | (Tatsuki et al. 2016) | mice | knockout | Increase sleep duration, 5.36min (1.07 SD) longer than WT. |
| II | *BDNF* | (Garner et al. 2018) | mice | knockdown(heterozygous) | Higher % of wakefulness, higher number of wake episodes, shorten duration of wake episodes,  less number of nREM episodes and shorten duration of nREM episodes. |
| II | *BLOC1S6* | (F. Y. Lee et al. 2018) | mice | knockout | Short sleep. |
| II | *CACNA1A* | (Deboer et al. 2013) | mice | R192Q knock-in | More waking with longer waking episodes in the dark period;  Less non-rapid eye movement (NERM) sleep but equal amounts of REM sleep. |
| II | *CACNA1B* | (Beuckmann et al. 2003) | mice | knockout | Normal total sleep duration;  Increased consolidation of REM sleep (less REM sleep episode);  Overall EEG spectral power increased during wakefulness and REM sleep, and decreased in NREM sleep. |
| II | *CACNA1G* | (J. Lee, Kim, and Shin 2004)., (Anderson et al. 2005). | mice | knockout | ~10% less NREM sleep;  Loss of the thalamic delta (1-4Hz) waves and reduction of sleep spindles (7-14Hz).  Slow rhythms(<1Hz) were relatively intact during urethane anesthesia and nREM sleep. |
| II | *CAMK2A* | (Tatsuki et al. 2016) | mice | knockout | Decrease in sleep duration, 50.1min (1.27SD) shorter than WT. |
| II | *CAMK2B* | (Tatsuki et al. 2016) | mice | knockout | Decrease in sleep duration, 120.3min (3.04SD) shorter than WT. |
| II | *CHRM1* | (Niwa et al. 2018) | mice | Chrm1/Chrm3 double knockout | Diminish REM sleep. |
| II | *CHRM3* | (Niwa et al. 2018) | mice | Chrm3-/- | Theta peak frequency for REM sleep was significantly increased. |
| II | *CLOCK* | (Naylor et al. 2000) | mice | homo-, hetero- Clock mutant | Reduce sleep during light periods, with reduction in the mean NREM episode length;  No difference between the number of brief arousals from sleep. |
| II | *CNTNAP2* | (Thomas et al. 2017a) | rat, mice | knockout | Reduced spectral power in the alpha range during wake;  Rat: increased consolidation of wakefulness and REM sleep;  Mice: wake fragmentation. |
| II | *CREB1* | (Graves et al. 2003) | mice | CREB αΔ homozygous | Decreased wakefulness;  Increased NREM sleep;  No change for average sleep bout length. |
| II | *CRH* | (Kimura et al. 2010) | mice | overexpression in whole brain or forebrain | Increase REM sleep;  Slightly suppressed nonREM sleep;  CRY hypersecretion in the forebrain seems to drive REM sleep. |
| II | *CSNK1E* | (Zhou et al. 2014) | mice | knockout | Higher proportion of sleep time spent in REM sleep during the dark period. |
| II | *DBH* | (Ouyang et al. 2004) | mice | knockout | Increase total sleep (~2hr/day);  Decrease the duration of waking bouts and increase number of waking bouts;  Decrease in REM sleep. |
| II | *DBP* | (P. Franken et al. 2000) | mice | knockout | Amplitude of circadian modulation of sleep time and consolidation of sleep episodes reduced;  Reduction in amplitude of sleep-wake-dependent changes in slow wave sleep delta power;  Increase in hippocampal theta peak frequency. |
| II | *EGR3* | (Maple et al. 2018) | mice | knockout | More wake-related behaviour. |
| II | *EIF4EBP1* | (Areal et al. 2020) | mice | knockout | Increase sleep duration and longer sleep bout (~23 second) during dark phase. |
| II | *FAAH* | (Huitron-Resendiz et al. 2004) | mice | knockout | <15% less waking and and <10% more NREM sleep during the light phase. |
| II | *FAH* | (Yang et al. 2019) | mice | N68S | Phase advanced and less total activity time in young mouse;  Earlier onset of activity (several hours before lights off);  A reduction in total activity and body weight when compared with wild-type or heterozygous mice;  These behavioral phenotypes became milder as the mice grew older and were completely rescued by the administration of NTBC. |
| II | *FMR1* | (Saré et al. 2017) | mice | fragile X syndrome model (dfmr1 and Fmr1 knockout/Fxr2 heterozygote) | Phenotype take place in adult (p70) animal but not in younger age;  Reduce total sleep duration in light phase. |
| II | *FOS* | (Shiromani et al. 2000) | mice | knockout | ~27% more waking/24hr (during sleep phase);  Reduce slow wave sleep  ~25% less NREM/24hr. |
| II | *FOSB* | (Shiromani et al. 2000) | mice | knockout | ~35% less REM/24hr. |
| II | *FUS* | (T. Zhang et al. 2018) | rat | R521C knock-in | Fragmented sleep and occurred during day and night phases;  REM decreases in R521C KI rats only in the first 6 hours in the dark phase. |
| II | *GRIA1* | (Ang et al. 2018) | mice | knockout | Reduction of EEG power density including the spindle-frequency range (10–15 Hz) during sleep;  Longer REM sleep episodes;  Increased EEG slow-wave activity in the occipital derivation during baseline sleep;  Reduced rate of decline of EEG slow wave activity (0.5–4 Hz) during NREM sleep after sleep deprivation. |
| II | *GRM2* | (Pritchett et al. 2015) | mice | Grm2/3 double knockout | Reduce sleep time;  Increase sleep fragmentation. |
| II | *GRM3* | (Pritchett et al. 2015) | mice | Grm2/3 double knockout | Reduce sleep time;  Increase sleep fragmentation |
| II | *HCRT* | (Vassalli and Franken 2017) | mice | knockout | Narcolepsy, drive sleep-to-wake transitions. |
| II | *HCRTR2* | (Lin et al. 1999; Chemelli et al. 1999; Hara et al. 2001) | canine, dog | autosomal recessive mutation | Narcolepsy |
| II | *HDC* | (Parmentier et al. 2002) | mice | knockout | Lower level of arousal  hypersomnolence;  ~23% more REM sleep/24h  decrease waking around the light to dark transition. |
| II | *HOMER1* | (Diering et al. 2017; Naidoo et al. 2012) | mice/drosophila | *homer* - R102 mutant drosophila, homer1a knockout mice | Drosophila - fragmented sleep and failure to sustain long bouts of sleep  Mice - failure to sustain long bouts of wakefulness. |
| II | *HTR1A* | (Benjamin Boutrel et al. 2002) | mice | knockout | ~50% more REM sleep/24hr. |
| II | *HTR1B* | (B. Boutrel et al. 1999) | mice | knockout | ~40% more REM sleep/24hr;  Lower amounts of slow-wave sleep during the light phase;  No REM sleep rebound after deprivation. |
| II | *HTR2A* | (Popa et al. 2005) | mice | knockout | Decrease NREM. |
| II | *HTR2C* | (Frank, Stryker, and Tecott 2002) | mice | knockout | ~20% more waking/24h;  ~20% less NREM sleep/24h;  Response to sleep deprivation: increase in NREM sleep duration, larger increase in delta activity. |
| II | *HTR7* | (Hedlund et al. 2005) | mice | knockout | Less time and less frequent episodes of REM sleep. |
| II | *IFNAR1* | (Bohnet et al. 2004) | mice | knockout | ~30% reduction in REM sleep;  No change for NREM sleep. |
| II | *IL1R1* | (Fang, Wang, and Krueger 1998) | mice | knockout | Decreased NREM during active time. |
| II | *IL6* | (Morrow and Opp 2005) | mice | knockout | Spend more time in NREM sleep after sleep deprivation. |
| II | *KCNA2* | (Douglas et al. 2007). | mice | knockout | Decrease NREM sleep;  Increase number of waking episodes;  No change in the number or duration of sleep. |
| II | *KCNC1* | (Espinosa et al. 2004) | mice | double mutant of Kcnc1/Kcnc3 | Show more significant phenotype  restlessness particularly prominent during light period (rest phase). |
| II | *KCNK9* | (Yoshida et al. 2018) | mice | knockout | Short sleep duration;  Decrease in sleep-state stabilization and increase in wake-state stabilization. |
| II | *KCNN3* | (Tatsuki et al. 2016) | mice | knockout | 106.6min(~2.1SD) shorter than WT;  Increase in awake-state stabilization, but not decrease in sleep-state stabilization. |
| II | *LEP* | (Laposky et al. 2006) | mice | male ob/ob mice | Elevated number of arousals from sleep;  Increased stage shifts;  More frequent, shorter-lasting sleep bouts - impair sleep consolidation  increase of NREM sleep/24hour. |
| II | *MCHR1* | (Adamantidis et al. 2008) | mice | Mch-R1 -/- | Normal total sleep duration / 24 hour  19% more REM sleep in the light phase. |
| II | *MEF2D* | (Mohawk et al. 2019) | mice | knockout in SCN | Phase delayed;  Longer free running period (40 minutes more than WT);  Greater number of activity bound per day;  More transition from NREM sleep to wakefulness and from wakefulness to NREM sleep;  Extreme fragmented sleep with significantly shorter REM latency. |
| II | *NALCN* | (Funato et al. 2016) | mice | random mutagenized | Reduce REM sleep. |
| II | *NFKB1* | (Jhaveri et al. 2006) | mice | knockout | Longer sleep duration  26% more sleep/24 hr;  More time in slow wave sleep (SWS) and rapid eye movement sleep (REMS) under normal conditions. |
| II | *NLGN2* | (Seok et al. 2018) | mice | knockout | More wakefulness, less NREM especially during dark period  higher absolute delta activity. |
| II | *NLGN3* | (Thomas et al. 2017b) | rat | knockout | Increase REM and decrease NREM time;  Elevated theta power(4-9Hz) during wakefulness and REM;  Elevated delta power(0.5-4Hz) during NREM;  Latency to enter NREM or REM sleep after light onset did not differ. |
| II | *NLRP3* | (Zielinski et al. 2017) | mice | knockout | Reduce NREM during light period. |
| II | *NOS1* | (L. Chen, Majde, and Krueger 2003) | mice | knockout | REM/nREM sleep ratio bidirectional alteration;  Neuronal nitric oxide synthase(nNOS) lowered REM sleep; inducible NOS increases REM sleep. |
| II | *NPAS2* | (Dudley et al. 2003; Paul Franken et al. 2006) | mice | knockout | Decrease NREM in second half of dark period;  Recovery of sleep time lost is impaired. |
| II | *NTSR1* | (Fitzpatrick et al. 2012) | mice | knockout | Short sleep;  Lower percentage of sleep time spent in REM sleep in the dark phase;  Larger diurnal variation in REM sleep duration than wild types under baseline conditions. |
| II | *OPN4* | (Tsai et al. 2009) | mice | knockout | Shorter sleep;  Light failed to induce sleep. |
| II | *PANX1* | (Kovalzon et al. 2017) | mice | knockout | Increase in waking;  Increase in movement activity;  Decrease in slow wave sleep percentages, especially during dark period. |
| II | *PER1* | (Kopp et al. 2002) | mice | knockout | No change in total sleep duration or sleep homeostasis;  Alter 24 hour sleep distribution. |
| II | *PRKG1* | (Langmesser et al. 2009) | mice | knockout in brain | No different in total sleep duration but more awake in the light phase;  ~25% decrease in REM sleep  reduced NREM delta activity during baseline. |
| II | *PRL* | (Obál et al. 2005) | mice | knockout | ~30% REM sleep reduction (light phase only). |
| II | *PROK2* | (Hu et al. 2007) | mice | knockout | Decrease in NREM sleep time;  Increase REM sleep time in both light and dark periods. |
| II | *PTPRD* | (Drgonova et al. 2015) | mice | hetero- and homo- knockout | 22% less sleep in homozygous knockout;  Restless leg syndrome. |
| II | *RAB3A* | (Kapfhamer et al. 2002) | mice | hD77G, resulting in reduced protein level | ~25% increase NREM selep/24h. |
| II | *RIMS1* | (Lonart et al. 2008) | mice | knockout | ~50% less baseline REM sleep;  ~35% less NREM sleep in dark phase only. |
| II | *SCN1A* | (Papale et al. 2013) | mice | human SCN1A R1648H mutants | Increase wakefulness;  Reduced non-rapid eye movement and rapid eye movement sleep during the dark phase. |
| II | *SCN8A* | (Papale et al. 2010) | mice | knockout | More nonREM sleep during dark phase;  Less REM sleep during light phase. |
| II | *SHANK3* | (Ingiosi et al. 2019) | mice | lack of exon 21 | Have problem falling asleep even if sleepy;  Less sleep during dark phase;  Alter EEG spectral under baseline. |
| II | *SIK3* | (Funato et al. 2016; Honda et al. 2018) | mice | random mutagenized  intron mutation cause skipping exon 13 mRNA  further validate that phosphorylation site S551 play the main role | Decrease in total wake time;  Increase NREM sleep;  Increase sleep need and normal wake promoting response. |
| II | *SIRT1* | (Panossian et al. 2011) | mice | hetero- and homo- knockout | Impaired wakefulness;  Decrease total wake time during active phase;  No disruption in REM/NREM sleep and sleep consolidation. |
| II | *SLC29A1* | (Kim et al. 2015) | mice | knockout | Decreased sleep duration;  Baseline NREM sleep decreases during the light period. |
| II | *SLC6A3* | (Wisor et al. 2001) | mice, canine | knockout | Increased wakefulness consolidation and promoting wakefulness;  Reduced nREM sleep. |
| II | *SLC6A4* | (Wisor et al. 2003) | mice | knockout | More frequent REMS bouts that last longer. |
| II | *TNF* | (Deboer, Fontana, and Tobler 2002) | mice | ligand knockout and TNFR2 knockout (R2KO) | Reduce REM sleep during the baseline light period due to a reduction in REM sleep episode frequency. |
| II | *TNFRSF1A* | (Fang, Wang, and Krueger 1997) | mice | knockout | ~20% less NREM sleep;  ~20% less REM sleep. |
| II | *UBE3A* | (Ehlen et al. 2015) | mice | Ube3am-/P+ | Sleep more in the early dark phase compare to control, lack of siesta during late dark phase;  More fragmented nighttime NREM sleep;  Reduce accumulation of sleep pressure. |
| II | *VAMP2* | (Banks et al. 2020) | mice | I102N | Shorter REM sleep;  Increased wakefulness and longer wake episodes. |
| III | *CUL3* | (Stavropoulos and Young 2011) | drosophila | Cul3 - knockdown | Reduced sleep duration, consolidation and homeostasis. |
| III | *ELP3* | (Singh et al. 2010) | drosophila | *Elp3* - knockout during development | Hyperactive phenotype and sleep loss in the adult flies. |
| III | *GRIN1* | (Tomita et al. 2015) | drosophila | *NMDAR1* - antagonist | Decrease total sleep;  Decrease sleep bout number. |
| III | *HTT* | (Faragó, Zsindely, and Bodai 2019) | drosophila | hHTTex1Q120, extra 95 glutamine at exon 1 | Reduce amount of sleep, sleep fragmentation, prolonged sleep latency. |
| III | *KCNA3* | (Cirelli et al. 2005) | drosophila | Screening - *Shaker* | ~60-85% decrease in daily sleep amount (mainly due to decrease in sleep episode duration, hyperactivity). |
| III | *KCTD5* | (Pfeiffenberger and Allada 2012) | drosophila | *Inc* - insertion | ~10h sleep reduction  hyper-arousability to a mechanical stimulus in adult flies. |
| III | *NEDD8* | (Stavropoulos and Young 2011) | drosophila | *Nedd8* - RNAi | Less total sleep time / 24 hour. |
| III | *PPP3CA* | (Tomita et al. 2011) | drosophila | *CanA14F* - Cn RNAi to knockdown the expression | Total sleep decrease to half or one-fifth of control;  Shorter sleep-bout length only during nighttime;  Sleep-bout number was reduced during the subjective day;  Overexpression induces sleep. |
| III | *PPP3R1* | (Tomita et al. 2011) | drosophila | *CanB* - Cn RNAi to knockdown the expression | Total sleep decrease to half or one-fifth of control;  Shorter sleep-bout length regardless of day or nighttime;  Sleep-bout number was reduced during the subjective day. |
| III | *PRKAB2* | (Nagy et al. 2018) | drosophila | *PRKAB2* - expressed alc-RNAi in nervous system | Reduced overall sleep, during both day and night phases;  Increase in duration of wakeful periods during light. |
| III | *RCAN2* | (Nakai et al. 2011) | drosophila | *Sra* - knockout | Two fold more active than WT;  ~30% less sleep than WT. |
| III | *SHMT1* | (Dai et al. 2019) | drosophila(intestine) | *SHMT* - knockout | Defective in D-ser synthesis: shorter sleep;  Defective in NMDAR1: shorter sleep;  Defective in D-ser degradation: longer sleep. |
| III | *SLC18A2* | (Nall and Sehgal 2013) | drosophila | *VMAT* - knockout | Increased arousal threshold. |
| III | *SLC6A2* | (Kume et al. 2005) | drosophila | *Fmn* - spontaneous mutation | Increased in total daily activity in LD and DD. |

Douglas, Christopher L., Vladyslav Vyazovskiy, Teresa Southard, Shing-Yan Chiu, Albee Messing, Giulio Tononi, and Chiara Cirelli. 2007. “Sleep in Kcna2 Knockout Mice.” *BMC Biology* 5 (October): 42.

Drgonova, Jana, Donna Walther, Katherine J. Wang, G. Luke Hartstein, Bryson Lochte, Juan Troncoso, Noriko Uetani, Yoichiro Iwakura, and George R. Uhl. 2015. “Mouse Model for Protein Tyrosine Phosphatase D (PTPRD) Associations with Restless Leg Syndrome or Willis-Ekbom Disease and Addiction: Reduced Expression Alters Locomotion, Sleep Behaviors and Cocaine-Conditioned Place Preference.” *Molecular Medicine*  21 (1): 717–25.

Dudley, Carol A., Claudia Erbel-Sieler, Sandi Jo Estill, Martin Reick, Paul Franken, Sinae Pitts, and Steven L. McKnight. 2003. “Altered Patterns of Sleep and Behavioral Adaptability in NPAS2-Deficient Mice.” *Science*  301 (5631): 379–83.

Ehlen, J. Christopher, Allison J. Brager, Julie Baggs, Lennisha Pinckney, Cloe L. Gray, Jason P. DeBruyne, Karyn A. Esser, Joseph S. Takahashi, and Ketema N. Paul. 2017. “Bmal1 Function in Skeletal Muscle Regulates Sleep.” *eLife* 6 (July). https://doi.org/[10.7554/eLife.26557](http://dx.doi.org/10.7554/eLife.26557).

Ehlen, J. Christopher, Kelly A. Jones, Lennisha Pinckney, Cloe L. Gray, Susan Burette, Richard J. Weinberg, Jennifer A. Evans, et al. 2015. “Maternal Ube3a Loss Disrupts Sleep Homeostasis But Leaves Circadian Rhythmicity Largely Intact.” *The Journal of Neuroscience: The Official Journal of the Society for Neuroscience* 35 (40): 13587–98.

Espinosa, F., G. Marks, N. Heintz, and R. H. Joho. 2004. “Increased Motor Drive and Sleep Loss in Mice Lacking Kv3-Type Potassium Channels.” *Genes, Brain, and Behavior* 3 (2): 90–100.

Fang, J., Y. Wang, and J. M. Krueger. 1997. “Mice Lacking the TNF 55 kDa Receptor Fail to Sleep More after TNFalpha Treatment.” *The Journal of Neuroscience: The Official Journal of the Society for Neuroscience* 17 (15): 5949–55.

———. 1998. “Effects of Interleukin-1 Beta on Sleep Are Mediated by the Type I Receptor.” *The American Journal of Physiology* 274 (3): R655–60.

Faragó, Anikó, Nóra Zsindely, and László Bodai. 2019. “Mutant Huntingtin Disturbs Circadian Clock Gene Expression and Sleep Patterns in Drosophila.” *Scientific Reports* 9 (1): 7174.

Fitzpatrick, Karrie, Christopher J. Winrow, Anthony L. Gotter, Joshua Millstein, Janna Arbuzova, Joseph Brunner, Andrew Kasarskis, Martha H. Vitaterna, John J. Renger, and Fred W. Turek. 2012. “Altered Sleep and Affect in the Neurotensin Receptor 1 Knockout Mouse.” *Sleep* 35 (7): 949–56.

Franken, Paul, Carol A. Dudley, Sandi Jo Estill, Monique Barakat, Ryan Thomason, Bruce F. O’Hara, and Steven L. McKnight. 2006. “NPAS2 as a Transcriptional Regulator of Non-Rapid Eye Movement Sleep: Genotype and Sex Interactions.” *Proceedings of the National Academy of Sciences of the United States of America* 103 (18): 7118–23.

Franken, P., L. Lopez-Molina, L. Marcacci, U. Schibler, and M. Tafti. 2000. “The Transcription Factor DBP Affects Circadian Sleep Consolidation and Rhythmic EEG Activity.” *The Journal of Neuroscience: The Official Journal of the Society for Neuroscience* 20 (2): 617–25.

Frank, Marcos G., Michael P. Stryker, and Laurence H. Tecott. 2002. “Sleep and Sleep Homeostasis in Mice Lacking the 5-HT2c Receptor.” *Neuropsychopharmacology: Official Publication of the American College of Neuropsychopharmacology* 27 (5): 869–73.

Funato, Hiromasa, Chika Miyoshi, Tomoyuki Fujiyama, Takeshi Kanda, Makito Sato, Zhiqiang Wang, Jing Ma, et al. 2016. “Forward-Genetics Analysis of Sleep in Randomly Mutagenized Mice.” *Nature* 539 (7629): 378–83.

Garner, Jennifer M., Jonathan Chambers, Abigail K. Barnes, and Subimal Datta. 2018. “Changes in Brain-Derived Neurotrophic Factor Expression Influence Sleep-Wake Activity and Homeostatic Regulation of Rapid Eye Movement Sleep.” *Sleep* 41 (2). https://doi.org/[10.1093/sleep/zsx194](http://dx.doi.org/10.1093/sleep/zsx194).

Gerstner, Jason R., Isaac J. Perron, Samantha M. Riedy, Takeo Yoshikawa, Hiroshi Kadotani, Yuji Owada, Hans P. A. Van Dongen, et al. 2017. “Normal Sleep Requires the Astrocyte Brain-Type Fatty Acid Binding Protein FABP7.” *Science Advances* 3 (4): e1602663.

Graves, Laurel A., Kevin Hellman, Sigrid Veasey, Julie A. Blendy, Allan I. Pack, and Ted Abel. 2003. “Genetic Evidence for a Role of CREB in Sustained Cortical Arousal.” *Journal of Neurophysiology* 90 (2): 1152–59.

Hara, J., C. T. Beuckmann, T. Nambu, J. T. Willie, R. M. Chemelli, C. M. Sinton, F. Sugiyama, et al. 2001. “Genetic Ablation of Orexin Neurons in Mice Results in Narcolepsy, Hypophagia, and Obesity.” *Neuron* 30 (2): 345–54.

Hedlund, Peter B., Salvador Huitron-Resendiz, Steven J. Henriksen, and J. Gregor Sutcliffe. 2005. “5-HT7 Receptor Inhibition and Inactivation Induce Antidepressantlike Behavior and Sleep Pattern.” *Biological Psychiatry* 58 (10): 831–37.

He, Ying, Christopher R. Jones, Nobuhiro Fujiki, Ying Xu, Bin Guo, Jimmy L. Holder Jr, Moritz J. Rossner, Seiji Nishino, and Ying-Hui Fu. 2009. “The Transcriptional Repressor DEC2 Regulates Sleep Length in Mammals.” *Science* 325 (5942): 866–70.

Hirano, Arisa, Guangsen Shi, Christopher R. Jones, Anna Lipzen, Len A. Pennacchio, Ying Xu, William C. Hallows, et al. 2016. “A Cryptochrome 2 Mutation Yields Advanced Sleep Phase in Humans.” *eLife* 5 (August). https://doi.org/[10.7554/eLife.16695](http://dx.doi.org/10.7554/eLife.16695).

Honda, Takato, Tomoyuki Fujiyama, Chika Miyoshi, Aya Ikkyu, Noriko Hotta-Hirashima, Satomi Kanno, Seiya Mizuno, et al. 2018. “A Single Phosphorylation Site of SIK3 Regulates Daily Sleep Amounts and Sleep Need in Mice.” *Proceedings of the National Academy of Sciences of the United States of America* 115 (41): 10458–63.

Huang, Zhi-Li, Wei-Min Qu, Naomi Eguchi, Jiang-Fan Chen, Michael A. Schwarzschild, Bertil B. Fredholm, Yoshihiro Urade, and Osamu Hayaishi. 2005. “Adenosine A2A, but Not A1, Receptors Mediate the Arousal Effect of Caffeine.” *Nature Neuroscience* 8 (7): 858–59.

Huitrón-Reséndiz, Salvador, Manuel Sánchez-Alavez, Roger Gallegos, Greta Berg, Elena Crawford, Jeannie L. Giacchino, Dora Games, Steven J. Henriksen, and José R. Criado. 2002. “Age-Independent and Age-Related Deficits in Visuospatial Learning, Sleep-Wake States, Thermoregulation and Motor Activity in PDAPP Mice.” *Brain Research* 928 (1-2): 126–37.

Huitron-Resendiz, Salvador, Manuel Sanchez-Alavez, Derek N. Wills, Benjamin F. Cravatt, and Steven J. Henriksen. 2004. “Characterization of the Sleep-Wake Patterns in Mice Lacking Fatty Acid Amide Hydrolase.” *Sleep* 27 (5): 857–65.

Hu, Wang-Ping, Jia-Da Li, Chengkang Zhang, Lisa Boehmer, Jerome M. Siegel, and Qun-Yong Zhou. 2007. “Altered Circadian and Homeostatic Sleep Regulation in Prokineticin 2-Deficient Mice.” *Sleep* 30 (3): 247–56.

Ingiosi, Ashley M., Hannah Schoch, Taylor Wintler, Kristan G. Singletary, Dario Righelli, Leandro G. Roser, Elizabeth Medina, Davide Risso, Marcos G. Frank, and Lucia Peixoto. 2019. “Shank3 Modulates Sleep and Expression of Circadian Transcription Factors.” *eLife* 8 (April). https://doi.org/[10.7554/eLife.42819](http://dx.doi.org/10.7554/eLife.42819).

Jaaro-Peled, Hanna, Cara Altimus, Tara LeGates, Tyler Cash-Padgett, Sandra Zoubovsky, Takatoshi Hikida, Koko Ishizuka, Samer Hattar, Valérie Mongrain, and Akira Sawa. 2016. “Abnormal Wake/sleep Pattern in a Novel Gain-of-Function Model of DISC1.” *Neuroscience Research* 112 (November): 63–69.

Jhaveri, K. A., V. Ramkumar, R. A. Trammell, and L. A. Toth. 2006. “Spontaneous, Homeostatic, and Inflammation-Induced Sleep in NF-κB p50 Knockout Mice.” *American Journal of Physiology-Regulatory, Integrative and Comparative Physiology*, November. https://doi.org/[10.1152/ajpregu.00262.2006](http://dx.doi.org/10.1152/ajpregu.00262.2006).

Kapfhamer, David, Otto Valladares, Yi Sun, Patrick M. Nolan, John J. Rux, Steven E. Arnold, Sigrid C. Veasey, and Maja Bućan. 2002. “Mutations in Rab3a Alter Circadian Period and Homeostatic Response to Sleep Loss in the Mouse.” *Nature Genetics* 32 (2): 290–95.

Kim, T., V. Ramesh, M. Dworak, D-S Choi, R. W. McCarley, A. V. Kalinchuk, and R. Basheer. 2015. “Disrupted Sleep-Wake Regulation in Type 1 Equilibrative Nucleoside Transporter Knockout Mice.” *Neuroscience* 303 (September): 211–19.

Kimura, M., P. Müller-Preuss, A. Lu, E. Wiesner, C. Flachskamm, W. Wurst, F. Holsboer, and J. M. Deussing. 2010. “Conditional Corticotropin-Releasing Hormone Overexpression in the Mouse Forebrain Enhances Rapid Eye Movement Sleep.” *Molecular Psychiatry* 15 (2): 154–65.

Kopp, Caroline, Urs Albrecht, Binhai Zheng, and Irene Tobler. 2002. “Homeostatic Sleep Regulation Is Preserved in mPer1 and mPer2 Mutant Mice.” *The European Journal of Neuroscience* 16 (6): 1099–1106.

Kovalzon, V. M., L. S. Moiseenko, A. V. Ambaryan, S. Kurtenbach, V. I. Shestopalov, and Y. V. Panchin. 2017. “Sleep-Wakefulness Cycle and Behavior in pannexin1 Knockout Mice.” *Behavioural Brain Research* 318 (February): 24–27.

Kume, Kazuhiko, Shoen Kume, Sang Ki Park, Jay Hirsh, and F. Rob Jackson. 2005. “Dopamine Is a Regulator of Arousal in the Fruit Fly.” *The Journal of Neuroscience: The Official Journal of the Society for Neuroscience* 25 (32): 7377–84.

Kurien, Philip, Pei-Ken Hsu, Jacy Leon, David Wu, Thomas McMahon, Guangsen Shi, Ying Xu, et al. 2019. “TIMELESS Mutation Alters Phase Responsiveness and Causes Advanced Sleep Phase.” *Proceedings of the National Academy of Sciences of the United States of America* 116 (24): 12045–53.

Langmesser, Sonja, Paul Franken, Susanne Feil, Yann Emmenegger, Urs Albrecht, and Robert Feil. 2009. “cGMP-Dependent Protein Kinase Type I Is Implicated in the Regulation of the Timing and Quality of Sleep and Wakefulness.” *PloS One* 4 (1): e4238.

Laposky, Aaron D., Jonathan Shelton, Joseph Bass, Christine Dugovic, Nicholas Perrino, and Fred W. Turek. 2006. “Altered Sleep Regulation in Leptin-Deficient Mice.” *American Journal of Physiology. Regulatory, Integrative and Comparative Physiology* 290 (4): R894–903.

Lee, Frank Y., Huei-Bin Wang, Olivia N. Hitchcock, Dawn Hsiao Loh, Daniel S. Whittaker, Yoon-Sik Kim, Achilles Aiken, et al. 2018. “Sleep/Wake Disruption in a Mouse Model of BLOC-1 Deficiency.” *Frontiers in Neuroscience* 12 (November): 759.

Lee, Jungryun, Daesoo Kim, and Hee-Sup Shin. 2004. “Lack of Delta Waves and Sleep Disturbances during Non-Rapid Eye Movement Sleep in Mice Lacking alpha1G-Subunit of T-Type Calcium Channels.” *Proceedings of the National Academy of Sciences of the United States of America* 101 (52): 18195–99.

Lin, L., J. Faraco, R. Li, H. Kadotani, W. Rogers, X. Lin, X. Qiu, P. J. de Jong, S. Nishino, and E. Mignot. 1999. “The Sleep Disorder Canine Narcolepsy Is Caused by a Mutation in the Hypocretin (orexin) Receptor 2 Gene.” *Cell* 98 (3): 365–76.

Lonart, G., X. Tang, F. Simsek-Duran, M. Machida, and L. D. Sanford. 2008. “The Role of Active Zone Protein Rab3 Interacting Molecule 1 Alpha in the Regulation of Norepinephrine Release, Response to Novelty, and Sleep.” *Neuroscience* 154 (2): 821–31.

Maple, Amanda M., Rachel K. Rowe, Jonathan Lifshitz, Fabian Fernandez, and Amelia L. Gallitano. 2018. “Influence of Schizophrenia-Associated Gene Egr3 on Sleep Behavior and Circadian Rhythms in Mice.” *Journal of Biological Rhythms* 33 (6): 662–70.

Medori, R., H. J. Tritschler, A. LeBlanc, F. Villare, V. Manetto, H. Y. Chen, R. Xue, S. Leal, P. Montagna, and P. Cortelli. 1992. “Fatal Familial Insomnia, a Prion Disease with a Mutation at Codon 178 of the Prion Protein Gene.” *The New England Journal of Medicine* 326 (7): 444–49.

Mohawk, Jennifer A., Kimberly H. Cox, Makito Sato, Seung-Hee Yoo, Masashi Yanagisawa, Eric N. Olson, and Joseph S. Takahashi. 2019. “Neuronal Myocyte-Specific Enhancer Factor 2D (MEF2D) Is Required for Normal Circadian and Sleep Behavior in Mice.” *The Journal of Neuroscience: The Official Journal of the Society for Neuroscience* 39 (40): 7958–67.

Morrow, Jonathan D., and Mark R. Opp. 2005. “Sleep-Wake Behavior and Responses of Interleukin-6-Deficient Mice to Sleep Deprivation.” *Brain, Behavior, and Immunity* 19 (1): 28–39.

Nagy, Stanislav, Gianna W. Maurer, Julie L. Hentze, Morten Rose, Thomas M. Werge, and Kim Rewitz. 2018. “AMPK Signaling Linked to the Schizophrenia-Associated 1q21.1 Deletion Is Required for Neuronal and Sleep Maintenance.” *PLoS Genetics* 14 (12): e1007623.

Naidoo, Nirinjini, Megan Ferber, Raymond J. Galante, Blake McShane, Jia Hua Hu, John Zimmerman, Greg Maislin, et al. 2012. “Role of Homer Proteins in the Maintenance of Sleep-Wake States.” *PloS One* 7 (4): e35174.

Nakai, Yasuhiro, Junjiro Horiuchi, Manabu Tsuda, Satomi Takeo, Shin Akahori, Takashi Matsuo, Kazuhiko Kume, and Toshiro Aigaki. 2011. “Calcineurin and Its Regulator sra/DSCR1 Are Essential for Sleep in Drosophila.” *The Journal of Neuroscience: The Official Journal of the Society for Neuroscience* 31 (36): 12759–66.

Nall, Aleksandra H., and Amita Sehgal. 2013. “Small-Molecule Screen in Adult Drosophila Identifies VMAT as a Regulator of Sleep.” *The Journal of Neuroscience: The Official Journal of the Society for Neuroscience* 33 (19): 8534–40.

Naylor, E., B. M. Bergmann, K. Krauski, P. C. Zee, J. S. Takahashi, M. H. Vitaterna, and F. W. Turek. 2000. “The Circadian Clock Mutation Alters Sleep Homeostasis in the Mouse.” *The Journal of Neuroscience: The Official Journal of the Society for Neuroscience* 20 (21): 8138–43.

Niwa, Yasutaka, Genki N. Kanda, Rikuhiro G. Yamada, Shoi Shi, Genshiro A. Sunagawa, Maki Ukai-Tadenuma, Hiroshi Fujishima, et al. 2018. “Muscarinic Acetylcholine Receptors Chrm1 and Chrm3 Are Essential for REM Sleep.” *Cell Reports* 24 (9): 2231–47.e7.

Obál, Ferenc, Jr, Fabio Garcia-Garcia, Balint Kacsóh, Ping Taishi, Stewart Bohnet, Nelson D. Horseman, and James M. Krueger. 2005. “Rapid Eye Movement Sleep Is Reduced in Prolactin-Deficient Mice.” *The Journal of Neuroscience: The Official Journal of the Society for Neuroscience* 25 (44): 10282–89.

Ouyang, Ming, Kevin Hellman, Ted Abel, and Steven A. Thomas. 2004. “Adrenergic Signaling Plays a Critical Role in the Maintenance of Waking and in the Regulation of REM Sleep.” *Journal of Neurophysiology* 92 (4): 2071–82.

Palchykova, Svitlana, Raphaelle Winsky-Sommerer, Hai-Ying Shen, Detlev Boison, Andrea Gerling, and Irene Tobler. 2010. “Manipulation of Adenosine Kinase Affects Sleep Regulation in Mice.” *The Journal of Neuroscience: The Official Journal of the Society for Neuroscience* 30 (39): 13157–65.

Panossian, Lori, Polina Fenik, Yan Zhu, Guanxia Zhan, Michael W. McBurney, and Sigrid Veasey. 2011. “SIRT1 Regulation of Wakefulness and Senescence-like Phenotype in Wake Neurons.” *The Journal of Neuroscience: The Official Journal of the Society for Neuroscience* 31 (11): 4025–36.

Papale, Ligia A., Christopher D. Makinson, J. Christopher Ehlen, Sergio Tufik, Michael J. Decker, Ketema N. Paul, and Andrew Escayg. 2013. “Altered Sleep Regulation in a Mouse Model of SCN1A-Derived Genetic Epilepsy with Febrile Seizures plus (GEFS+).” *Epilepsia* 54 (4): 625–34.

Papale, Ligia A., Ketema N. Paul, Nikki T. Sawyer, Joseph R. Manns, Sergio Tufik, and Andrew Escayg. 2010. “Dysfunction of the Scn8a Voltage-Gated Sodium Channel Alters Sleep Architecture, Reduces Diurnal Corticosterone Levels, and Enhances Spatial Memory.” *The Journal of Biological Chemistry* 285 (22): 16553–61.

Parmentier, Régis, Hiroshi Ohtsu, Zahia Djebbara-Hannas, Jean-Louis Valatx, Takehiko Watanabe, and Jian-Sheng Lin. 2002. “Anatomical, Physiological, and Pharmacological Characteristics of Histidine Decarboxylase Knock-out Mice: Evidence for the Role of Brain Histamine in Behavioral and Sleep-Wake Control.” *The Journal of Neuroscience: The Official Journal of the Society for Neuroscience* 22 (17): 7695–7711.

Patke, Alina, Patricia J. Murphy, Onur Emre Onat, Ana C. Krieger, Tayfun Özçelik, Scott S. Campbell, and Michael W. Young. 2017. “Mutation of the Human Circadian Clock Gene CRY1 in Familial Delayed Sleep Phase Disorder.” *Cell* 169 (2): 203–15.e13.

Pfeiffenberger, Cory, and Ravi Allada. 2012. “Cul3 and the BTB Adaptor Insomniac Are Key Regulators of Sleep Homeostasis and a Dopamine Arousal Pathway in Drosophila.” *PLoS Genetics* 8 (10): e1003003.

Popa, Daniela, Clément Léna, Véronique Fabre, Caroline Prenat, Jay Gingrich, Pierre Escourrou, Michel Hamon, and Joëlle Adrien. 2005. “Contribution of 5-HT2 Receptor Subtypes to Sleep-Wakefulness and Respiratory Control, and Functional Adaptations in Knock-out Mice Lacking 5-HT2A Receptors.” *The Journal of Neuroscience: The Official Journal of the Society for Neuroscience* 25 (49): 11231–38.

Pritchett, David, Aarti Jagannath, Laurence A. Brown, Shu K. E. Tam, Sibah Hasan, Silvia Gatti, Paul J. Harrison, David M. Bannerman, Russell G. Foster, and Stuart N. Peirson. 2015. “Deletion of Metabotropic Glutamate Receptors 2 and 3 (mGlu2 & mGlu3) in Mice Disrupts Sleep and Wheel-Running Activity, and Increases the Sensitivity of the Circadian System to Light.” *PloS One* 10 (5): e0125523.

Saré, R. Michelle, Lee Harkless, Merlin Levine, Anita Torossian, Carrie A. Sheeler, and Carolyn B. Smith. 2017. “Deficient Sleep in Mouse Models of Fragile X Syndrome.” *Frontiers in Molecular Neuroscience* 10 (September): 280.

Seok, Bong Soo, Feng Cao, Erika Bélanger-Nelson, Chloé Provost, Steve Gibbs, Zhengping Jia, and Valérie Mongrain. 2018. “The Effect of Neuroligin-2 Absence on Sleep Architecture and Electroencephalographic Activity in Mice.” *Molecular Brain* 11 (1): 52.

Shi, Guangsen, Lijuan Xing, David Wu, Bula J. Bhattacharyya, Christopher R. Jones, Thomas McMahon, S. Y. Christin Chong, et al. 2019. “A Rare Mutation of β1-Adrenergic Receptor Affects Sleep/Wake Behaviors.” *Neuron* 0 (0). https://doi.org/[10.1016/j.neuron.2019.07.026](http://dx.doi.org/10.1016/j.neuron.2019.07.026).

Shi, Guangsen, Chen Yin, Zenghua Fan, Lijuan Xing, Yulia Mostovoy, Pui-Yan Kwok, Liza H. Ashbrook, Andrew D. Krystal, Louis J. Ptáček, and Ying-Hui Fu. 2020. “Mutations in Metabotropic Glutamate Receptor 1 Contribute to Natural Short Sleep Trait.” *Current Biology: CB*, October. https://doi.org/[10.1016/j.cub.2020.09.071](http://dx.doi.org/10.1016/j.cub.2020.09.071).

Shiromani, P. J., R. Basheer, J. Thakkar, D. Wagner, M. A. Greco, and M. E. Charness. 2000. “Sleep and Wakefulness in c-Fos and Fos B Gene Knockout Mice.” *Brain Research. Molecular Brain Research* 80 (1): 75–87.

Singh, Neetu, Meridith T. Lorbeck, Ashley Zervos, John Zimmerman, and Felice Elefant. 2010. “The Histone Acetyltransferase Elp3 Plays in Active Role in the Control of Synaptic Bouton Expansion and Sleep in Drosophila.” *Journal of Neurochemistry* 115 (2): 493–504.

Stavropoulos, Nicholas, and Michael W. Young. 2011. “Insomniac and Cullin-3 Regulate Sleep and Wakefulness in Drosophila.” *Neuron* 72 (6): 964–76.

Tatsuki, Fumiya, Genshiro A. Sunagawa, Shoi Shi, Etsuo A. Susaki, Hiroko Yukinaga, Dimitri Perrin, Kenta Sumiyama, et al. 2016. “Involvement of Ca(2+)-Dependent Hyperpolarization in Sleep Duration in Mammals.” *Neuron* 90 (1): 70–85.

Thomas, Alexia M., Michael D. Schwartz, Michael D. Saxe, and Thomas S. Kilduff. 2017a. “Cntnap2 Knockout Rats and Mice Exhibit Epileptiform Activity and Abnormal Sleep-Wake Physiology.” *Sleep* 40 (1). https://doi.org/[10.1093/sleep/zsw026](http://dx.doi.org/10.1093/sleep/zsw026).

———. 2017b. “Sleep/Wake Physiology and Quantitative Electroencephalogram Analysis of the Neuroligin-3 Knockout Rat Model of Autism Spectrum Disorder.” *Sleep* 40 (10). https://doi.org/[10.1093/sleep/zsx138](http://dx.doi.org/10.1093/sleep/zsx138).

Tobler, I., S. E. Gaus, T. Deboer, P. Achermann, M. Fischer, T. Rülicke, M. Moser, B. Oesch, P. A. McBride, and J. C. Manson. 1996. “Altered Circadian Activity Rhythms and Sleep in Mice Devoid of Prion Protein.” *Nature* 380 (6575): 639–42.

Tomita, Jun, Madoka Mitsuyoshi, Taro Ueno, Yoshinori Aso, Hiromu Tanimoto, Yasuhiro Nakai, Toshiro Aigaki, Shoen Kume, and Kazuhiko Kume. 2011. “Pan-Neuronal Knockdown of Calcineurin Reduces Sleep in the Fruit Fly, Drosophila Melanogaster.” *The Journal of Neuroscience: The Official Journal of the Society for Neuroscience* 31 (37): 13137–46.

Tomita, Jun, Taro Ueno, Madoka Mitsuyoshi, Shoen Kume, and Kazuhiko Kume. 2015. “The NMDA Receptor Promotes Sleep in the Fruit Fly, Drosophila Melanogaster.” *PloS One* 10 (5): e0128101.

Tsai, Jessica W., Jens Hannibal, Grace Hagiwara, Damien Colas, Elisabeth Ruppert, Norman F. Ruby, H. Craig Heller, Paul Franken, and Patrice Bourgin. 2009. “Melanopsin as a Sleep Modulator: Circadian Gating of the Direct Effects of Light on Sleep and Altered Sleep Homeostasis in Opn4(-/-) Mice.” *PLoS Biology* 7 (6): e1000125.

Vanselow, Katja, Jens T. Vanselow, Pål O. Westermark, Silke Reischl, Bert Maier, Thomas Korte, Andreas Herrmann, Hanspeter Herzel, Andreas Schlosser, and Achim Kramer. 2006. “Differential Effects of PER2 Phosphorylation: Molecular Basis for the Human Familial Advanced Sleep Phase Syndrome (FASPS).” *Genes & Development* 20 (19): 2660–72.

Vassalli, Anne, and Paul Franken. 2017. “Hypocretin (orexin) Is Critical in Sustaining Theta/gamma-Rich Waking Behaviors That Drive Sleep Need.” *Proceedings of the National Academy of Sciences of the United States of America* 114 (27): E5464–73.

Wisor, J. P., S. Nishino, I. Sora, G. H. Uhl, E. Mignot, and D. M. Edgar. 2001. “Dopaminergic Role in Stimulant-Induced Wakefulness.” *The Journal of Neuroscience: The Official Journal of the Society for Neuroscience* 21 (5): 1787–94.

Wisor, J. P., S. W. Wurts, F. S. Hall, K. P. Lesch, D. L. Murphy, G. R. Uhl, and D. M. Edgar. 2003. “Altered Rapid Eye Movement Sleep Timing in Serotonin Transporter Knockout Mice.” *Neuroreport* 14 (2): 233–38.

Xing, Lijuan, Guangsen Shi, Yulia Mostovoy, Nicholas W. Gentry, Zenghua Fan, Thomas B. McMahon, Pui-Yan Kwok, Christopher R. Jones, Louis J. Ptáček, and Ying-Hui Fu. 2019. “Mutant Neuropeptide S Receptor Reduces Sleep Duration with Preserved Memory Consolidation.” *Science Translational Medicine* 11 (514). https://doi.org/[10.1126/scitranslmed.aax2014](http://dx.doi.org/10.1126/scitranslmed.aax2014).

Xu, Ying, Quasar S. Padiath, Robert E. Shapiro, Christopher R. Jones, Susan C. Wu, Noriko Saigoh, Kazumasa Saigoh, Louis J. Ptácek, and Ying-Hui Fu. 2005. “Functional Consequences of a CKIdelta Mutation Causing Familial Advanced Sleep Phase Syndrome.” *Nature* 434 (7033): 640–44.

Yang, Shuzhang, Sandra M. Siepka, Kimberly H. Cox, Vivek Kumar, Marleen de Groot, Yogarany Chelliah, Jun Chen, Benjamin Tu, and Joseph S. Takahashi. 2019. “Tissue-Specific FAH Deficiency Alters Sleep-Wake Patterns and Results in Chronic Tyrosinemia in Mice.” *Proceedings of the National Academy of Sciences of the United States of America* 116 (44): 22229–36.

Yoshida, Kensuke, Shoi Shi, Maki Ukai-Tadenuma, Hiroshi Fujishima, Rei-Ichiro Ohno, and Hiroki R. Ueda. 2018. “Leak Potassium Channels Regulate Sleep Duration.” *Proceedings of the National Academy of Sciences of the United States of America* 115 (40): E9459–68.

Zhang, Luoying, Arisa Hirano, Pei-Ken Hsu, Christopher R. Jones, Noriaki Sakai, Masashi Okuro, Thomas McMahon, et al. 2016. “A PERIOD3 Variant Causes a Circadian Phenotype and Is Associated with a Seasonal Mood Trait.” *Proceedings of the National Academy of Sciences of the United States of America* 113 (11): E1536–44.

Zhang, Tao, Xin Jiang, Min Xu, Haifang Wang, Xiao Sang, Meiling Qin, Puhua Bao, et al. 2018. “Sleep and Circadian Abnormalities Precede Cognitive Deficits in R521C FUS Knockin Rats.” *Neurobiology of Aging* 72 (December): 159–70.

Zhou, Lili, Camron D. Bryant, Andrew Loudon, Abraham A. Palmer, Martha Hotz Vitaterna, and Fred W. Turek. 2014. “The Circadian Clock Gene Csnk1e Regulates Rapid Eye Movement Sleep Amount, and Nonrapid Eye Movement Sleep Architecture in Mice.” *Sleep* 37 (4): 785–93, 793A – 793C.

Zielinski, Mark R., Dmitry Gerashchenko, Svetlana A. Karpova, Varun Konanki, Robert W. McCarley, Fayyaz S. Sutterwala, Robert E. Strecker, and Radhika Basheer. 2017. “The NLRP3 Inflammasome Modulates Sleep and NREM Sleep Delta Power Induced by Spontaneous Wakefulness, Sleep Deprivation and Lipopolysaccharide.” *Brain, Behavior, and Immunity* 62 (May): 137–50.
